# Resolving the climatic and ecological drivers of geographical gradients in avian sexual selection

**DOI:** 10.1101/2023.09.01.555923

**Authors:** Robert A. Barber, Jingyi Yang, Chenyue Yang, Oonagh Barker, Tim Janicke, Joseph A. Tobias

## Abstract

Sexual selection, one of the central pillars of evolutionary theory, has powerful effects on organismal morphology, behaviour and population dynamics. However, current knowledge about geographical variation in this evolutionary mechanism and its underlying drivers remains highly incomplete, in part because standardized data on the strength of sexual selection is sparse even for well-studied organisms. Here we use information on mating systems – including the incidence of polygamy and extra-pair paternity – to quantify the intensity of sexual selection in 10671 (>99.9%) bird species distributed worldwide. We show that avian sexual selection varies latitudinally, peaking at higher latitudes, although the gradient is reversed in the world’s most sexually selected birds – specialist frugivores – which are strongly associated with tropical forests. Phylogenetic models further reveal that the strength of sexual selection is explained by temperature seasonality coupled with a suite of climate-associated factors, including migration, diet, and territoriality. Overall, these analyses suggest that climatic conditions leading to short, intense breeding seasons, or highly abundant and patchy food resources, increase the potential for polygamy in birds, driving latitudinal gradients in sexual selection. Our findings help to resolve longstanding debates about spatial variation in evolutionary mechanisms linked to reproductive biology, and provide a comprehensive species-level dataset for further studies of selection and phenotypic evolution in the context of global climatic change.

## Introduction

Many of the most spectacular outcomes of animal evolution, from the peacock’s tail to the song of the nightingale, are testament to the power of sexual selection, a pervasive mechanism driven by competition among individuals for reproductive success (Andersson 1994). Decades of intensive research have shown that sexual selection has far-reaching impacts on adaptation (Bonduriansky 2011), speciation (Mendelson & Safran 2021), and population dynamics (Lumley *et al*. 2015), with implications for genomic evolution (Wyer *et al*. 2023) and the response of organisms to environmental change (Chenoweth *et al*. 2015; Parrett *et al*. 2019; Moiron *et al*. 2022). Previous studies focusing on local contexts or experimental systems have made substantial progress in understanding how sexual selection varies within populations (Twiss *et al*. 2006; Morimoto *et al*. 2016; Hare & Simmons 2022), yet the factors explaining variation among species remain unclear, particularly at wider taxonomic and geographic scales (Cornwallis & Uller 2010; Macías-Ordóñez *et al*. 2013; Miller & Svensson 2014; Machado *et al*. 2016).

Large-scale patterns in reproductive strategies have often been proposed, with one recurring suggestion being that sexual selection may vary with latitude or climate. However, both the existence and the direction of these potential gradients remain disputed (Fig. 1). Some studies suggest that sexual selection is likely to increase at higher latitudes where shorter reproductive periods and higher breeding synchrony intensify competition for mates (Catchpole 1980; Irwin 2000), and present more opportunities for polygyny (Emlen & Oring 1977) and extra-pair matings (Stutchbury & Morton 1995). Conversely, other studies suggest that sexual selection will peak in the tropics, either because year-round reproduction and asynchronous breeding allows males to mate with multiple females sequentially (Møller & Ninni 1998), or because long fruiting or flowering seasons can promote either resource defence polygyny (Machado *et al*. 2016) or female-only parental care, leading to extreme forms of male-male competition, such as lekking (Beehler 1983; Barve & La Sorte 2016).

**Fig. 1.**
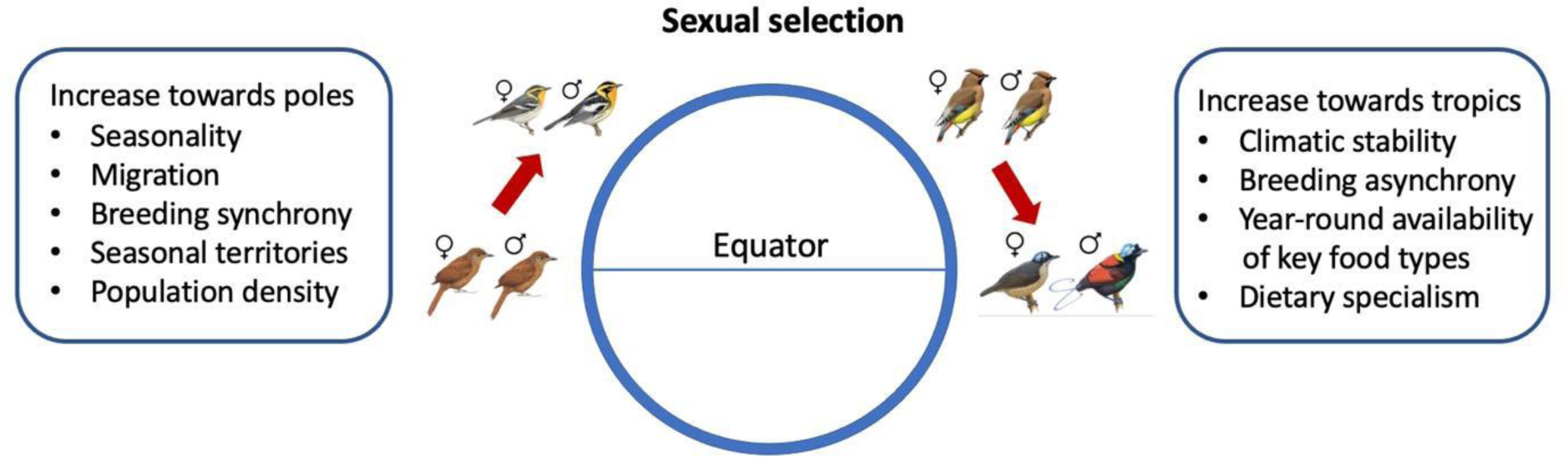
Hypothetical gradients in sexual selection and their putative mechanisms. One hypothesis proposes that avian sexual selection increases with latitude because highly seasonal climates promote short, synchronous breeding seasons, leading to intense competition for mating opportunities in dense populations. This concept may apply primarily to higher trophic levels, with a gradient from monomorphic insectivores with year-round territories in the tropics (illustrated by Uniform Treehunter) to highly dimorphic migratory insectivores at higher latitudes (for example, Blackburnian Warbler). An alternative hypothesis predicts that sexual selection increases towards the tropics because climatic stability promotes year-round availability of food resources, including abundant fruit and flowers, leading to extreme polygamy in dietary specialists. This concept may apply primarily to lower trophic levels, with a gradient from monomorphic frugivores with biparental care at higher latitudes (for example, Japanese Waxwing) to highly dimorphic frugivores with female-only care in the tropics (for example, Wilson’s Bird-of-Paradise).

These alternative hypotheses are difficult to disentangle, and previous tests have produced conflicting results. Some analyses suggest that sexual selection is most intense at higher latitudes (Irwin 2000; Mahler & Gil 2009; Weir & Wheatcroft 2011), whereas others find evidence that sexual selection either increases in the tropics (Barve & La Sorte 2016; Murray *et al*. 2021) or has no significant relationship with latitude in either direction (Cardillo 2002; Hendry *et al*. 2014; Friedman & Remeš 2016; Lifjeld *et al*. 2019). A major limitation is that numerous attempts to examine macroecological patterns in reproductive strategies have relied on secondary sexual traits, such as sexual size dimorphism or sexual dichromatism, as indices of sexual selection (Irwin 2000; Weir & Wheatcroft 2011; Svensson & Waller 2013; Dale *et al*. 2015; Murray *et al*. 2021; Pincheira-Donoso *et al*. 2021; Cooney *et al*. 2022). These metrics can only provide an incomplete picture of variation in reproductive strategies because they overlook sexual selection acting on different traits, such as acoustic signals (Cooney *et al*. 2018). The predictive power of standard morphological metrics for sexual selection are also weakened by mutual sexual selection (Kraaijeveld *et al*. 2007) and social selection (West-Eberhard 1979), both of which can drive increased body size and ornamentation in females (Tobias *et al*. 2012). A more complete test of sexual selection gradients and their drivers requires more direct evidence from behavioural observations and parentage analyses, yet this information has not previously been available at the required taxonomic and geographic scale.

To provide a new perspective on global gradients, we quantified variation in the strength of sexual selection inferred from mating behaviour across 10671 (>99.9%) bird species worldwide. These standardized scores of sexual selection were compiled using a well-established protocol based on the degree of polygamy (Verner & Willson 1966; Møller 1986; Owens & Hartley 1998; Dale *et al*. 2015) with modifications based on information about extra-pair paternity (EPP) and operational sex ratios (OSR). Although mating systems and associated metrics do not provide a complete measure of sexual selection (Jennions *et al*. 2012), they can serve as a robust and objective proxy (Krakauer *et al*. 2011; Janicke *et al*. 2016; Janicke & Morrow 2018) with fewer limitations than widely used morphological indices (Cooney *et al*. 2018).

Birds currently offer the best available system for a global synthesis of sexual selection because we know more about their breeding behaviour than any other major taxonomic group (Jetz & Rubenstein 2011; Valcu *et al*. 2021; Gonzalez-Voyer *et al*. 2022). In particular, a growing number of studies report information on relatively cryptic forms of sexual selection, including molecular analyses of parentage revealing the degree of EPP (Brouwer & Griffith 2019; Valcu *et al*. 2021). Furthermore, information on breeding can be coupled with uniquely comprehensive datasets on avian phylogenetic relationships, geographical distribution, life history and ecology (Jetz *et al*. 2012; Tobias *et al*. 2020, 2022). By combining these resources for all birds, along with GIS-based mapping and Bayesian models, we test key hypotheses about fundamental drivers of macroevolutionary patterns in sexual selection (Cornwallis & Uller 2010; Macías-Ordóñez *et al*. 2013).

We begin by assessing whether sexual selection scores are correlated with the latitude at which species occur, and then – because latitudinal gradients are best viewed as emergent properties shaped by multiple underlying mechanisms (Machado *et al*. 2016) – we evaluate the role of climate and ecology as fundamental drivers of variation in sexual selection across species. Specifically, we conduct a range of analyses to assess the relative importance of diet (Snow 1971; Beehler 1983), migratory behaviour (García-Peña *et al*. 2009), territoriality (Davies & Lundberg 1984) and climatic seasonality (Saenz *et al*. 2006), all of which have been proposed to regulate the intensity of sexual selection and thus to explain the enormous variation in sexual traits observed worldwide (Barve & La Sorte 2016; Machado *et al*. 2016; Tobias *et al*. 2016). Most hypotheses predict that sexual selection is accentuated by abundant food supply, migration, smaller home ranges, or seasonal environments, but the relevance of these factors at macroecological scales remains unknown.

## Results

We scored sexual selection by assigning 10671 bird species (Clements *et al*. 2021), including all 9988 valid bird species included in the global BirdTree phylogeny (Extended Data Fig. 1a) (Jetz *et al*. 2012), to one of five categories ranging from 0 (strict monogamy) to 4 (extreme polygamy). These scores were assigned according to thresholds based on the degree of polygamy, extra-pair paternity (EPP) or operational sex ratio (OSR); see Methods. Most BirdTree species (*n* = 8271; 83%) were scored as strictly monogamous, with higher levels of sexual selection (scores 1–4) assigned to the remainder (*n* = 1721, 17%; see Extended Data Fig. 1). The predominance of strict monogamy may partly reflect a tendency to underestimate sexual selection in the literature. For example, species observed in socially monogamous pairs are generally assumed to be genetically monogamous, whereas further study often reveals them to have higher rates of polygamy and EPP (Brouwer & Griffith 2019). Accordingly, when we focused exclusively on the best-known BirdTree species with highest certainty data (*n* = 2883), the proportion of strictly monogamous species was lower (*n* = 1937, 67%), while a third of species (n = 946, 33%) were classified as either polygamous or subject to sexual selection via EPP or OSR (scores 1–4). Sexual selection and data-certainty scores for all bird species, aligned with both Clements and BirdTree taxonomy, are presented along with sources in Supplementary Dataset 1.

To assess the validity of our sexual selection scores, we compared them against three alternative metrics of sexual selection: residual testes mass as an index of post-copulatory sperm competition (Dunn *et al*. 2001), Bateman gradients reflecting the increase in reproductive success obtained from additional matings (Bateman 1948), and opportunity for sexual selection (*I_S_*) reflecting variance in mating success (Wade 1979; for a full explanation, sources and derivation of these metrics see Supplementary Information). Bayesian phylogenetic models revealed significant positive relationships between our scores and all three metrics (Extended Data Fig. 2; Table S1). The amount of variation explained was relatively low for testes mass (*R*^2^ = 0.30), stronger for Bateman gradients (*R*^2^ = 0.53), and very strong for *I_S_* (*R*^2^ = 0.97) which provides a more accurate index of sexual selection (see Discussion).

### Geographical gradients of sexual selection

We overlaid geographical range data to map sexual selection scores for 9836 bird species (excluding species lacking spatial data), revealing lower average levels of sexual selection in tropical regions (Fig. 2a). Bayesian regression models (species level) confirmed that sexual selection follows a latitudinal gradient, increasing significantly towards higher latitudes (Fig. 2b). However, when we used the same procedure to assess geographic biases in knowledge, we found that average data certainty also increased in the temperate zone, with sexual selection scores being particularly robust in North America and Europe, where many bird species have been studied intensively for decades (Fig. 2c,d).

**Fig. 2.**
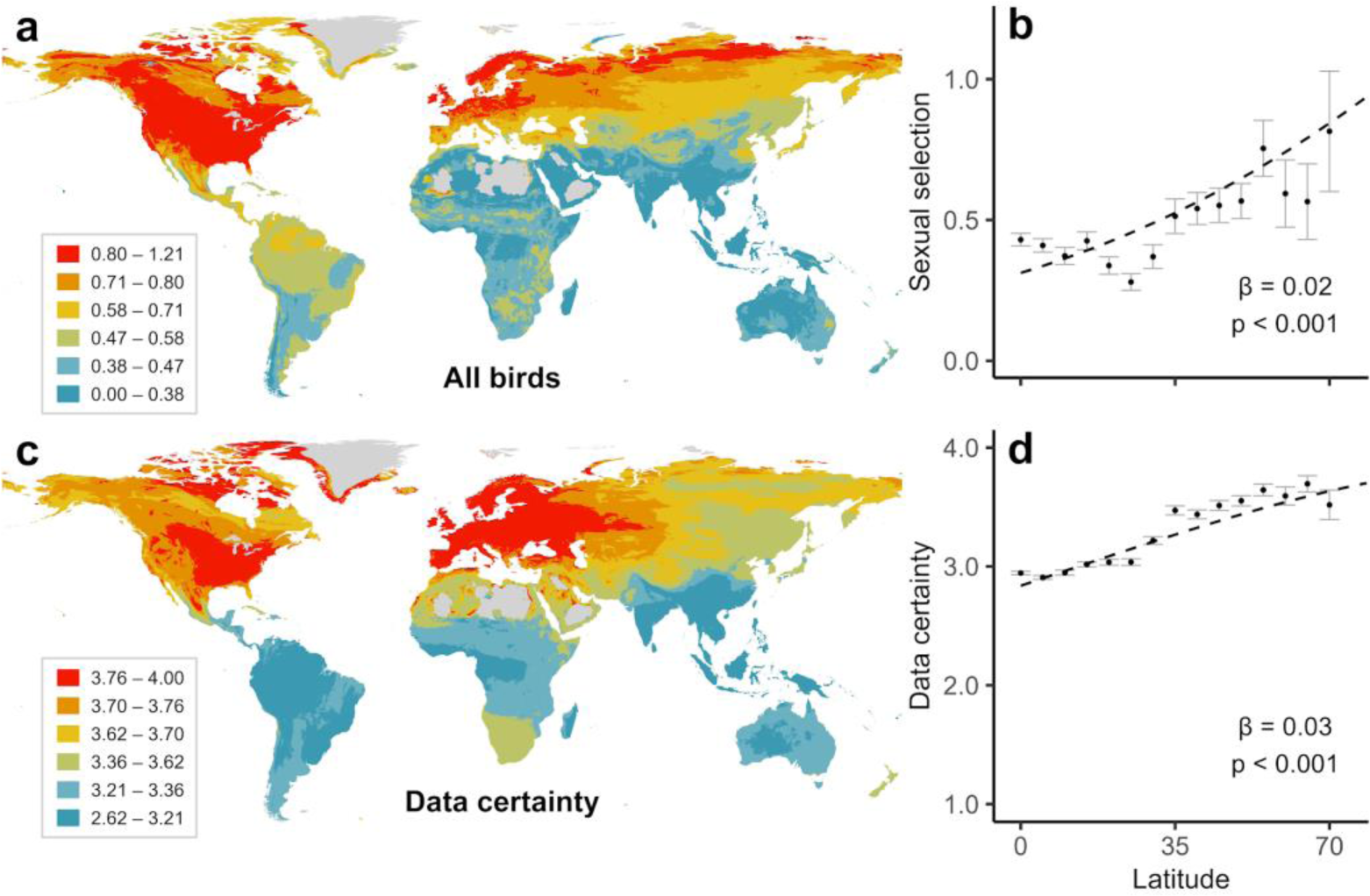
Geographical distribution of sexual selection in birds. **a**, Worldwide variation in sexual selection scores for 9836 bird species included in a global phylogeny (www.birdtree.org; Jetz *et al*. 2012), averaged for all species occurring in 5-km grid cells based on geographical range maps. Sexual selection was scored based on mating behaviour from monogamy (scored 0) to extreme polygamy (scored 4; see Extended Data Table 1). **b**, Relationship between avian sexual selection and latitude. The positive latitudinal gradient in sexual selection may seem counter-intuitive given the occurrence of largely monogamous clades such as seabirds and raptors at higher latitudes, but reflects intense sexual selection in other temperate and polar species, including both passerines (Verner & Wilson 1969) and non-passerines (Lanctot *et al*. 1997; Kempenaers & Valcu 2017; Küpper *et al*. 2016). **c**, Geographical variation in data certainty, ranging from no evidence (scored 1) to strong evidence (scored 4; see Extended Data Table 2). **d**, Relationship between data certainty and latitude. In **b** and **d**, variables were averaged within five degree latitudinal bins based on the centroid latitude of species’ geographical ranges; points represent means; bars denote one standard error; dashed lines were generated from species-level Bayesian regression models predicting sexual selection strength. Additional species-level models on a high certainty subset (scored 3–4) showed similar patterns (Table S2; Supplementary Information). In **a** and **c**, cells with <10 species were excluded from maps; in **b** and **d**, and latitudinal bins above 70 degrees were excluded from all scatterplots.

These patterns reflect a clear correlation between improved data quality and higher sexual selection scores (Fig. S1), suggesting that geographical variation in ornithological knowledge could potentially explain the overall latitudinal gradient in sexual selection. This does not appear to be the case, however, because the gradient in sexual selection remains strongly significant when we restrict the analysis to 7592 species with moderate to high data certainty (scored 3–4; Extended Data Fig. 3a,b).

Even when we restricted the analysis to the highest category of certainty (*n* = 2851 species), the positive relationship between sexual selection and latitude remained, albeit weaker because highly polygamous tropical species such as cotingas and manakins are more easily classified with certainty than monogamous species, explaining the accumulation of high-certainty species in the tropics (Fig. S2) and the relatively high average sexual selection score among the best-known Amazonian birds (Extended Data Fig. 3c,d).

### Univariate models predicting sexual selection

To understand the mechanisms driving latitudinal gradients in sexual selection, we re-examined geographical patterns through the lens of ecological traits. This approach revealed contrasting gradients when comparing primary consumers (herbivores, frugivores, granivores, and nectarivores) with secondary consumers (omnivores, carnivores, and scavengers). The positive latitudinal gradient in sexual selection is removed (non-significant) in primary consumers (Fig. 3a,b) and strongly reversed (negative) in an ecologically important subset: frugivores (Fig. 3c,d). Conversely, we found a strong positive gradient in secondary consumers (Fig. 3e,f), particularly the invertivores, which comprise ∼40% of the world’s birds (Fig. 3g,h). Thus, the overall positive gradient across all birds is predominantly driven by the species-rich invertivores, concealing an opposite (negative) gradient in fruit- and berry-eating species (frugivores), which constitute a smaller proportion (∼10%) of global bird diversity.

**Fig 3.**
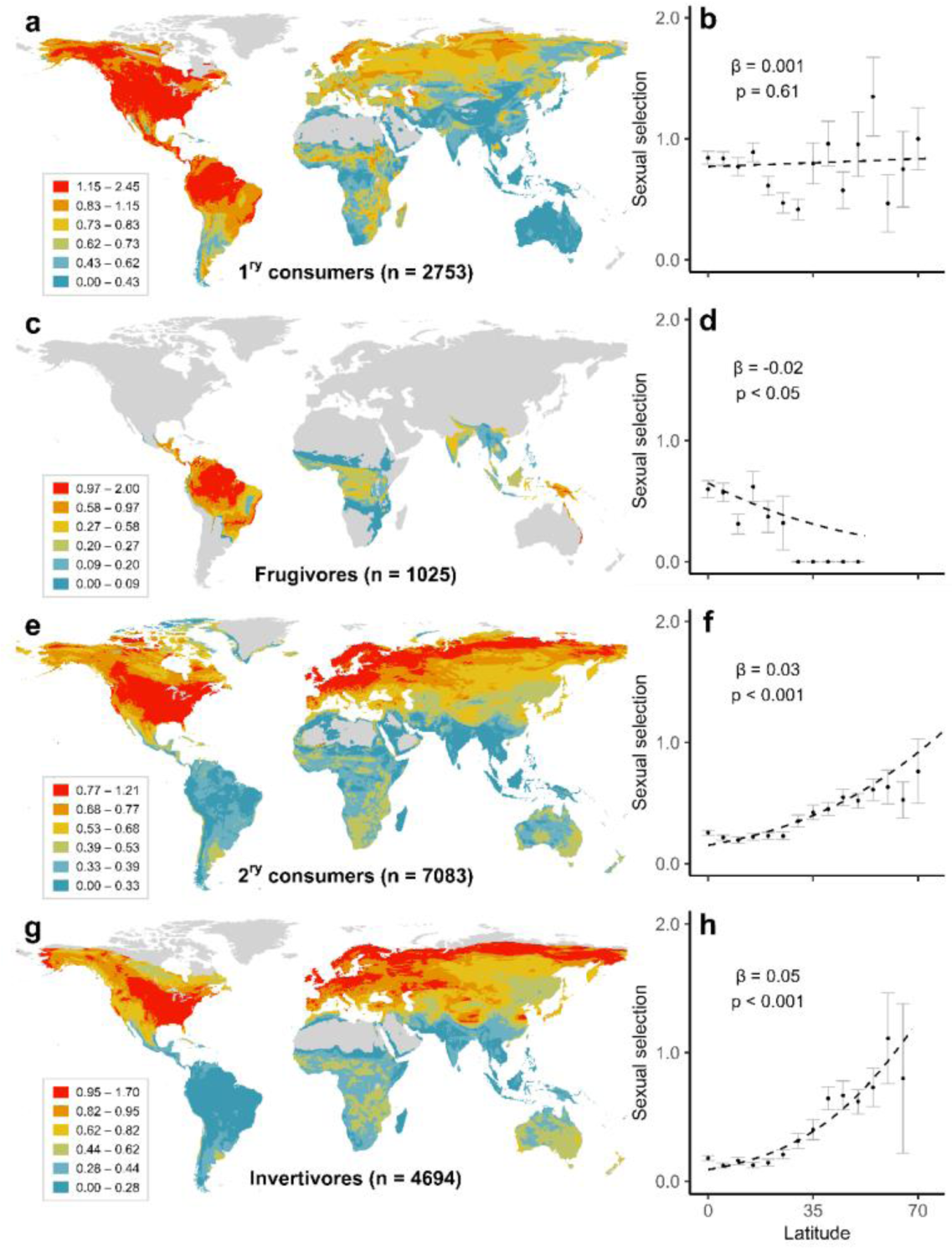
Global distribution of sexual selection partitioned by trophic level and diet. Upper panels show strength of sexual selection in primary (1^ry^) consumers mapped globally (**a**) and plotted against latitude (**b**), as well as for frugivores only (**c**,**d**). Lower panels show strength of sexual selection in secondary (2^ry^) consumers mapped globally (**e**) and plotted against latitude (**f**), as well as for invertivores only (**g**,**h**). We treated omnivores and carnivores as secondary consumers. Sexual selection was scored from monogamy (0) to extreme polygamy (4; see Methods). For mapping, we averaged sexual selection scores in each 5-km grid cell based on all species with geographical ranges overlapping the cell. In scatterplots, variables were averaged in 5-degree bins based on the centroid latitude of their ranges; points represent means; bars denote one standard error; dashed lines were generated from species-level Bayesian regression models. Sensitivity analyses on a subset of species with high-certainty data (scored 3–4) showed similar patterns (Table S2). To reduce noise, cells with <10 species were excluded from all maps, and latitudinal bins above 70 degrees were excluded from all scatterplots.

Switching focus to another key life history trait, migration, we found that both migratory and non-migratory species showed a significant positive gradient in sexual selection across latitude (Extended Data Fig. 4), aligning with the general trend observed across all birds. This consistent alignment was disrupted, however, when we focused on territoriality. While a strong positive gradient, twice the strength of the overall global relationship, was found in territorial species, the reverse pattern was found in non-territorial species, with substantially higher sexual selection observed in tropical regions (Extended Data Fig. 5). Similar variation in sexual selection across latitude in the context of ecological traits persisted when we classified bird species as either tropical or non-tropical (Extended Data Fig. 6). In all cases, the patterns were similar when we restricted analyses to well-known species (Table S2), and irrespective of how we quantified latitude (Table S3), indicating that our results are not biased by data certainty. These highly consistent results suggest that a combination of ecological strategies relating to diet, migration, and resource defence shape global trends in avian sexual selection. However, the relative importance of each trait is difficult to determine based on raw patterns given our finding that sexual selection is phylogenetically conserved in birds (Fig. 4a).

**Fig. 4.**
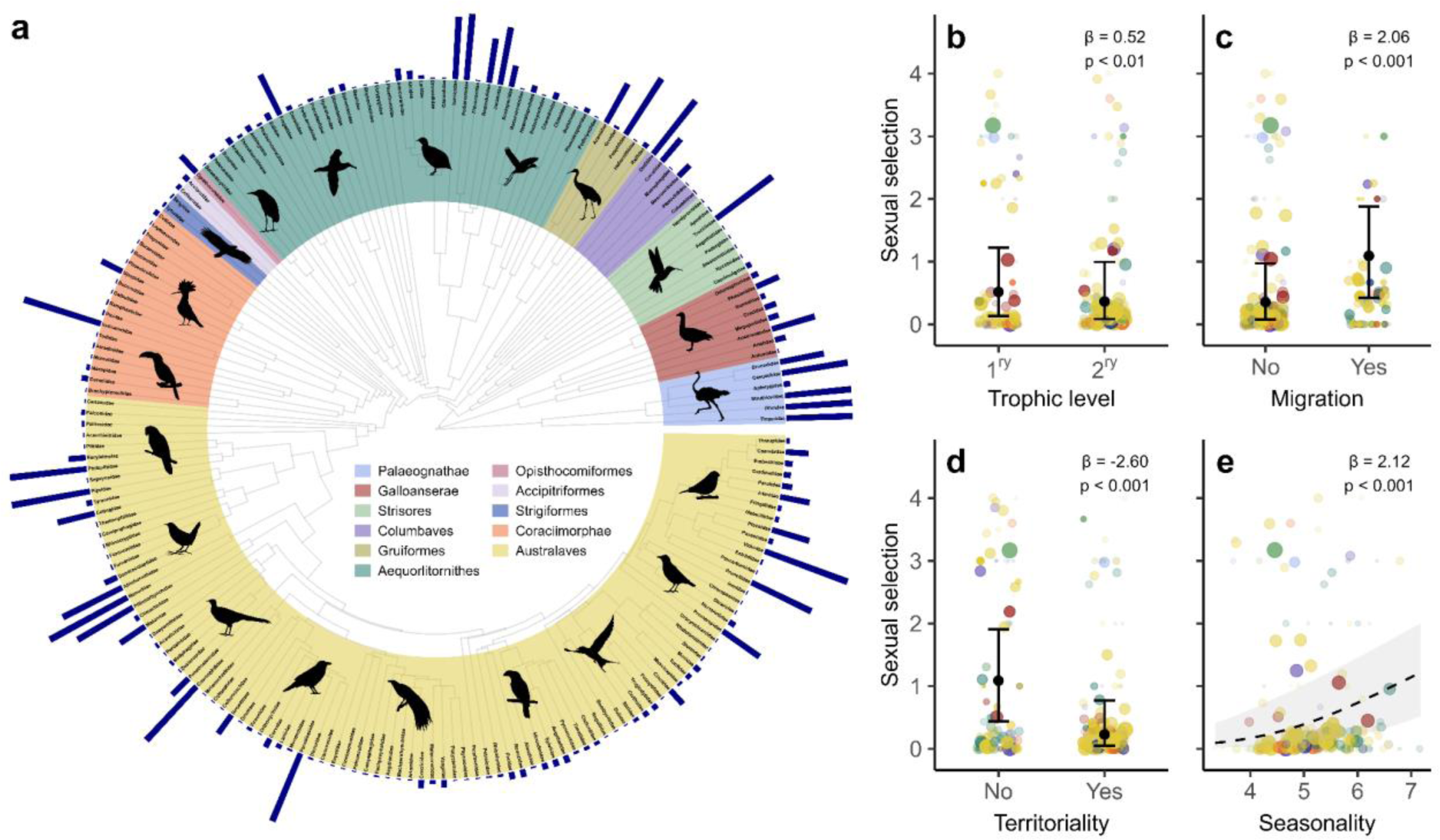
Potential role of climate in driving patterns of variation in sexual selection in birds. **a**, A family-level consensus phylogenetic tree (see Methods) showing the distribution of sexual selection across 194 families (*n* = 9988 species). Bars at branch tips show relative strength of sexual selection, with longer bars denoting higher average scores (see Methods). Coloured segments demarcate major clades (named in legend); silhouettes depict representative families (sourced from phylopic.org). Dot-and-whisker plots (**b–d**) show average sexual selection scores for each family, partitioned by trophic level (**b**), migratory behaviour (**c**) and territoriality (**d**); black points show predicted mean values for each group from species-level Bayesian phylogenetic models; whiskers show 95% confidence intervals. In **b**, 1^ry^ = primary; 2^ry^ = secondary. Each species trait shown in **b**–**d** is hypothetically linked to an overriding climatic driver, temperature seasonality, which we also found to be independently related to levels of sexual selection (**e**); dashed line shows model fit from a species-level Bayesian phylogenetic model; shaded area denotes 95% confidence intervals from model predictions. In **b**–**e**, data points for each family are coloured by major clades, following the same colour scheme as in the phylogeny, with size and transparency of each data point scaled to within-family species richness. Statistics show effect size and p-value comparing main effects with reference groups: secondary consumer, no migration, and no territoriality, respectively. Additional species-level models on a high certainty subset (scored 3–4) showed similar patterns (Table S5).

Using Bayesian phylogenetic models to account for evolutionary non-independence, we re-assessed the correlation between ecological traits and sexual selection across all species. As a first step, we ran a species-level model including trophic level as a binary predictor, with phylogenetic co-variance as a random effect (see Methods). We then repeated the same univariate model for migration and territoriality. All three ecological traits emerged as significant predictors (Table S4; Table S5), with the intensity of sexual selection increasing in primary consumers (Fig. 4b), migratory species (Fig. 4c), and non-territorial species (Fig. 4d). These ecological traits are all strongly tied to climatic seasonality (Tobias *et al*. 2016; Sheard *et al*. 2020), which we also found is strongly associated with global trends in sexual selection score (Fig. 4e; Table S4; Table S5). This raises two important questions: first, whether any particular ecological trait is driving the overall patterns, and second, whether climatic factors play an important role, either directly or via their effect on species ecology.

### Multivariate models

To examine the relative roles of ecological and climatic factors, we included all predictors in a multivariate Bayesian phylogenetic model (Fig. 5; Table S4) along with two key interactions accounting for complex relationships between diet and territoriality (Tobias *et al*. 2016) and between diet and seasonality (Murphy *et al*. 2023). We found that the strongest constraint (negative effect) on sexual selection was territoriality, with significantly lower levels of sexual selection in species defending seasonal or year-round territories. Conversely, the strongest driver (positive effect) was climatic seasonality, with significantly higher levels of sexual selection in species breeding in the most variable climates. While accounting for both these effects, we found that trophic level and migration also retained their independent significant positive effects on avian sexual selection (Fig. 5; Table S4).

**Fig. 5.**
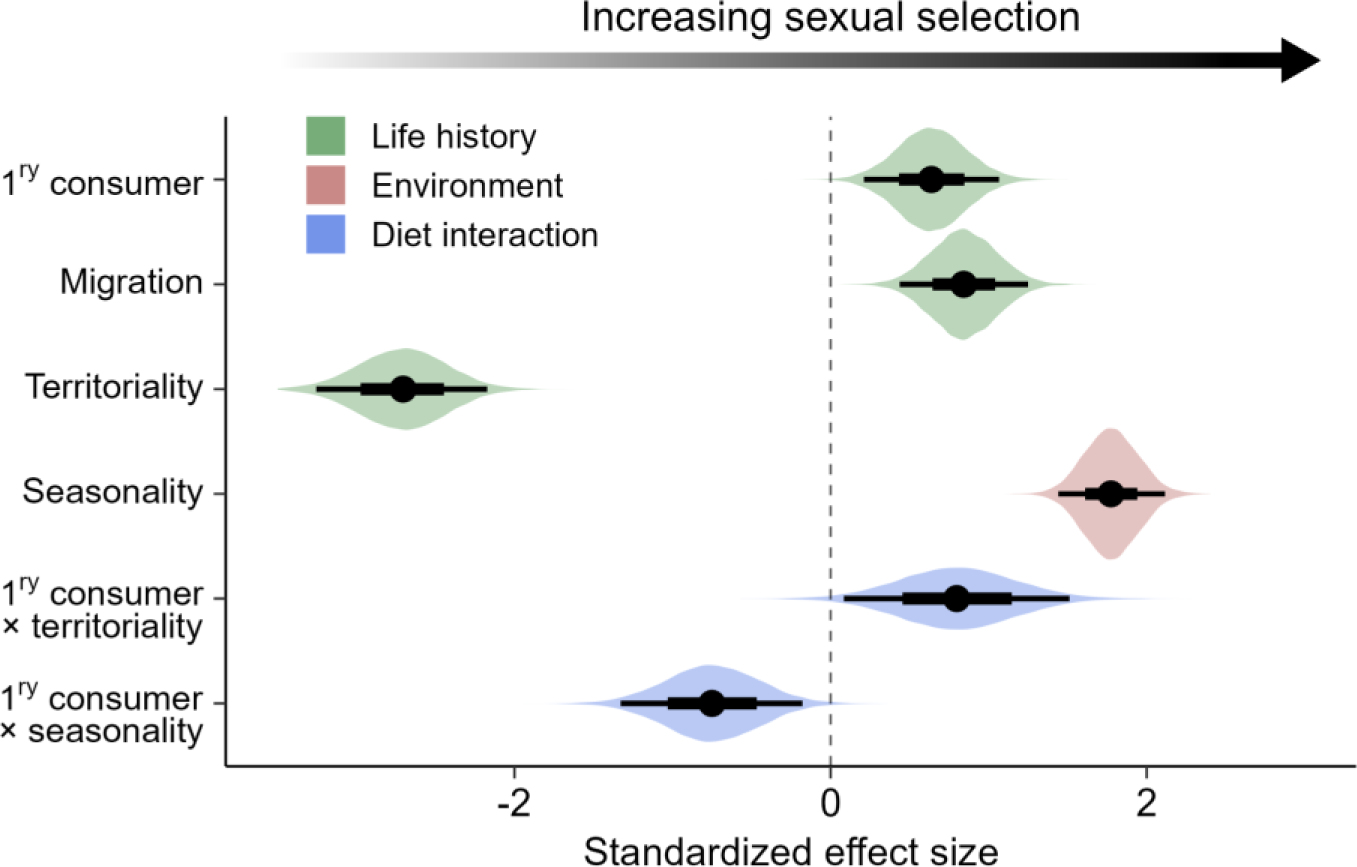
Relative roles of ecology and climate as drivers of sexual selection in birds. Results shown are from Bayesian phylogenetic models testing drivers of sexual selection in 9836 species. Predictors include three life history variables (green), one climatic variable (pink), and two key interactions between diet and the dominant effects (territoriality and seasonality; blue). 1ry consumer = primary consumer. The reference groups for the three categorical predictors are as follows: secondary consumer; no migration; and no territoriality, respectively (see Methods for definitions). Models were run on a sample of 50 phylogenetic trees extracted from www.birdtree.org (Jetz *et al*. 2012), grafted to the Prum *et al*. (2015) genomic backbone. Dots show mean effect size estimates from 12,500 posterior draws. For each effect, broad bases of whiskers show 66% confidence intervals; narrow tips of whiskers show 95% confidence intervals. Coloured distributions indicate the spread of effect size estimates, generated from a sample of 1000 posterior draws. Full statistical results are presented in Supplementary Table S4.

Although the independent effect of trophic level on sexual selection was relatively small, this ecological trait appears to have a key role in shaping variation in sexual selection among species through significant interactions with both territoriality (β = 0.80, 95% CI = 0.08, 1.51) and climatic seasonality (β = -0.75, 95% CI = -1.33, -0.18; Fig. 5; Table S4). In other words, dietary factors regulate the effects of territoriality and seasonality, which in turn have different implications for species at different trophic levels. We found that a phylogenetic model incorporating all ecological traits, along with the effects of seasonality and the key interactions with diet, explained a very high proportion of variation in sexual selection (*R*^2^ = 0.90; 95% CI = 0.88, 0.92). The proportion of variance explained became even stronger (*R*^2^ = 0.92; 95% CI = 0.91, 0.94) when we ran a more conservative analysis focusing exclusively on species with high-certainty data (Extended Data Fig. 7; Table S5).

## Discussion

Based on new estimates for all birds, our analyses confirm that sexual selection is subject to opposing latitudinal patterns, with the intensity of selection increasing towards higher latitudes in secondary consumers, contrasting with flat or reversed gradients identified in primary consumers (Fig. 1). These findings explain why positive latitudinal gradients of sexual selection tend to be proposed by authors studying insectivorous birds (Emlen & Oring 1977; Catchpole 1980; Irwin 2000) whereas negative gradients are proposed in studies of frugivores (Beehler 1983; Barve & La Sorte 2016). They may also help to resolve conflicting reports in the literature from a wider range of animal groups, with putative latitudinal gradients in sexual selection varying from positive (Saenz *et al*. 2006; Svensson & Waller 2013) to negative (Machado *et al*. 2016; Krasnov *et al*. 2022), as well as many studies reporting a lack of significant gradients when averaging across all species (Cardillo 2002; Hendry *et al*. 2014; Friedman & Remeš 2016; Lifjeld *et al*. 2019).

To unravel the mechanistic drivers of these complex and inconsistent global trends, we ran phylogenetic models showing that variation in sexual selection across birds is predominantly shaped by two interlinked factors – climatic seasonality and territoriality – both of which vary strongly with latitude. The strongest positive predictor of sexual selection across all birds was seasonality, in line with previous studies identifying a link between sexual selection and climatic heterogeneity, including variation in temperature (Pincheira-Donoso *et al*. 2021), rainfall (Saenz *et al*. 2006), or even local weather patterns (Twiss *et al*. 2006). An important consequence of climatic heterogeneity, including seasonality, is that key resources such as food and therefore reproductive individuals tend to cluster in time. As a result, species experience more intense competition for available mates during a shorter reproductive window (Stutchbury & Morton 2000), as well as a concentrated availability of mates and more efficient multiple matings (Stutchbury & Morton 1995). However, our findings clarify that climatic seasonality does not act alone, but in concert with other factors that can regulate or even reverse geographic gradients in sexual selection.

Among the ecological traits assessed in our analyses, territoriality was the strongest predictor of sexual selection, with a strong negative effect. This may be because territorial species tend to evolve monogamous mating systems (Komers & Brotherton 1997) or because the defence of non-overlapping territories automatically constrains population density and therefore limits access to additional mates and opportunistic matings. Davies & Lundberg (1984) illustrated this point by manipulating resource density in the territories of female Dunnocks (*Prunella modularis*), showing reduced territorial size and higher rates of polygamy as access to resources increased. Territories can have different implications for nectarivores, in which males of some species defend patches of flowers to attract potential mates (Temeles & Kress 2010).

Similarly, in mammalian systems, variation in social dominance can lead to variation in territory quality among males (Jarman 1974), and consequently increased mating success for those with the best territories (known as resource defence polygyny). While this may cause a positive link between territoriality and sexual selection in some avian systems, we find that the overall global pattern is reversed, presumably because resource defence polygyny is relatively rare.

To examine how the effects of climatic seasonality and resource defence strategies are influenced by diet more generally, we included interactions with trophic niches in our phylogenetic models. We found significant interactions with both seasonality and territoriality, indicating that trophic ecology plays a crucial role in shaping geographical variation in sexual selection. On the one hand, climatically stable environments such as tropical rainforests may be associated with increased sexual selection in primary consumers because the year-round abundance of nutrient-rich fruit and flowers favours breeding systems in which males are freed from the constraints of territory defence and parental care (leading to extreme polygyny in lekking frugivores and nectarivores). On the other hand, in secondary consumers such as insectivores, climatic stability promotes stable year-round pair- or group-territoriality (see Extended Data Fig. 8), explaining the steep gradient towards low levels of sexual selection in rainforest regions. These stable territorial systems are largely absent at temperate latitudes or in polar regions, where most insectivores are subject to much higher levels of sexual selection on ephemeral breeding territories before migrating to lower latitudes during winter (Stutchbury & Morton 2000; Winger *et al*. 2019).

We included migration in our models because migratory behaviour correlates with sexually selected traits and breeding systems across a range of taxa (Spottiswoode & Møller 2004; Albrecht *et al*. 2007; Dale *et al*. 2015; García-Roa *et al*. 2020). Even within single species, female choice has been shown to exert stronger sexual selection in migratory than sedentary populations (Collins *et al*. 2009). Despite the widespread consensus that migration promotes sexual selection, the effects can mostly be ascribed to climatic seasonality, which is very strongly correlated with migratory behaviour in birds (Somveille *et al*. 2015). Indeed, seasonal climatic fluctuations are the fundamental reason for the near-absence of insect prey, flowers and fruit during the winter period at higher latitudes, forcing many species to migrate (Winger *et al*. 2019). Intriguingly, our models reveal that, even after accounting for the effect of seasonality, migration remains weakly but significantly associated with sexual selection. This independent effect of migration is difficult to explain, perhaps relating to higher population density or more extreme breeding synchrony than resident species living in highly seasonal environments.

### The future of sexual selection in the context of climate change

Our findings indicate that geographical variation in sexual selection may be sensitive to shifts in seasonality projected to occur worldwide, and particularly in mid- to high latitudes (Xu *et al*. 2013). In mid-latitudes, reduced climatic fluctuations and milder winters may lead to increased survival of resident birds, in parallel with year-round availability of insects, nectar and fruit. This can reduce selection for migration (Winger *et al*. 2018), causing migratory bird populations to become sedentary (Pulido & Berthold 2010), while also increasing the duration of breeding seasons for resident species (Halupka & Halupka 2017). Our results suggest that these behavioural changes will dampen sexual selection, in line with previous studies showing that higher temperatures (García-Roa *et al*. 2020) and aseasonal climates (Botero & Rubenstein 2012; Ventura *et al*. 2021) can modify competition for mates and mating opportunities. Nonetheless, the precise impacts of climate change on mechanisms of sexual selection and their adaptive implications (Slatyer *et al*. 2012; Lumley *et al*. 2015; Parrett & Knell 2018; Wanders *et al*. 2023) are hard to predict because lengthening warm seasons have variable and species-specific effects on reproductive phenology in both plants (Linderholm 2006) and animals (Halupka & Halupka 2017; Harvey *et al*. 2023).

### Sexual selection scores for all birds: validity, robustness and research priorities

Our analyses offer the most comprehensive synthesis of geographical variation in sexual selection, estimated for all 9988 valid bird species aligned with a global phylogeny (Jetz *et al*. 2012). We provide this information on sexual selection, data sources and data certainty for all species in Supplementary Dataset 1, representing a major step-change in data availability from the most recent compilations of sexual selection scores, covering 3250 passerine species (Dale *et al*. 2015; Gonzalez-Voyer *et al*. 2022). We extended these earlier datasets, not only in terms of species coverage, but by scoring sexual selection based on a wider accumulation of evidence, including molecular evidence of polygamy and extra-pair paternity.

We found that these revised scores correlated relatively weakly with one well-established metric of sexual selection in birds: residual testes mass (Extended Data Figure S3). This was partly because residual testes mass appears to decline in species with the highest levels of sexual selection. This makes sense because females tend not remate in lekking systems (Avery 1984) and males consequently invest heavily in pre-copulatory sexual competition (Höglund & Alatalo 1995). Thus, testes mass reflects sperm competition but is a relatively crude metric for sexual selection. In contrast, our scores correlated strongly with two more sophisticated metrics – Bateman gradient and *I_S_* (Extended Data Figure S3) – suggesting that our data accurately reflect variance in mating success among individuals, at least in cases where sufficient evidence is available.

Our results also reveal a strong latitudinal gradient in data certainty (Fig. 2c,d), reflecting a long-standing bias towards ornithological research in the temperate zone (Stutchbury & Morton 2000) as well as the difficulty of tracking down relevant literature published in languages other than English. However, we note that most high-certainty species in our dataset are found in the tropics (Fig. S2), suggesting that biases in knowledge to some extent reflect the sheer number of tropical species compared with higher latitudes. The availability of high-certainty scores across all latitudes and ecological categories is crucial, allowing us to re-run our analyses with a more conservative dataset, confirming that all our main results are robust to biases in knowledge.

We hope that the open release of comprehensive estimates of sexual selection for over 10,000 bird species provides a template for further macro-scale research. Hypotheses linking sexual selection to evolutionary mechanisms and ecological contexts (Tobias & Seddon 2009; Biagolini-Jr. *et al*. 2017) can now be examined at unprecedented scale. We also hope our global dataset can re-energize efforts to improve the quality of evidence. In particular, the distribution of species with low-certainty data (Fig. 2c & S1) offers a detailed map of research priorities, highlighting the urgent need for observational and molecular studies, especially in the tropics, to quantify polygamy, EPP and OSR in species with low to moderate levels of sexual selection.

## Conclusions

Our analyses show that global-scale gradients in avian sexual selection are driven largely by temperature seasonality, with the strength of selection also strongly constrained by territoriality, promoted by migration, and modified by interactions with dietary niche. This complex but consistent interplay of environmental factors and species traits provides a clear explanation for the conflicting latitudinal patterns reported in previous studies, thus helping to resolve a long-standing debate about the validity of gradients in sexual selection and their underlying mechanisms (Macías-Ordóñez *et al*. 2013; Machado *et al*. 2016). These findings offer important insights into the origins and future of biodiversity, both in clarifying the fundamental eco-evolutionary processes driving global variation in mating systems and sexual traits (Cornwallis & Uller 2010) and providing a framework for understanding the evolutionary impacts of climatic change (Pulido & Berthold 2010; Marske *et al*. 2023).

## Methods

### Quantifying sexual selection

We scored sexual selection for all bird species using information published in primary and secondary literature, building on previous estimates of sexual selection (Dunn *et al*. 2001; Pitcher *et al*. 2005; Dale *et al*. 2015; Gonzalez-Voyer *et al*. 2022). Most information at family- and species-level was extracted from regional or global handbooks, including the Handbook of the Birds of the World series (del Hoyo *et al*. 1992-2013) with recent updates (Billerman *et al*. 2022). When published information was inconclusive, we used expert knowledge and personal observations to guide scoring decisions.

To assign scores, we adapted a well-established system based on the estimated proportion of polygamous individuals of either sex in the population (Verner & Willson 1966; Møller 1986; Owens & Hartley 1998; Dale *et al*. 2015). In addition, we further developed previous approaches by integrating information on EPP, OSR and display behaviour where possible. We assumed that high rates of EPP are associated with larger variance in male reproductive success (Møller & Ninni 1998), and thus reflect elevated levels of sexual selection even when species are socially monogamous. We also assumed that highly skewed OSR can intensify sexual selection (Janicke & Morrow 2018). Assignments were made based on quantitative estimates or textual descriptions of observed polygamy, extra-pair behaviour or sex ratios, with the addition of genetic estimates of EPP data, now available for a growing number of species (Brouwer & Griffith 2019).

Using the full set of criteria described in Extended Data Table 1, we assigned sexual selection scores of 0 (strict monogamy), 1 (frequent monogamy), 2 (regular polygamy), 3 (frequent polygamy) or 4 (extreme polygamy) to all species. We assumed that sexual selection increases from 0 to 4. Allocation to the first four categories was defined by well-established thresholds for polygamy (<0.1%, 0.1–5%, 5–20% and >20% for scores of 0–3, respectively). We used similar but higher quantitative thresholds for EPP data for two reasons. First, species may be strictly monogamous yet still have reported EPP rates above zero because of factors such as mate switching or brood parasitism (Pinxten *et al*. 1993). Consequently, following previous authors (Griffith *et al*. 2002), we define monogamy as <5% EPP. Second, many socially monogamous species have relatively high levels of EPP, so we adjusted thresholds upwards (assigning populations with 5–25%, 25–50% and >50% EPP to sexual selection scores 1–3, respectively). This avoids inflating sexual selection scores for species with EPP data, a biased subset of well-studied birds mainly inhabiting the temperate zone (Brouwer & Griffith 2019). Finally, species with highly elaborate mating display behaviours, including social lekking (e.g. Birds-of-Paradise, Paradisaeidae) or permanent display locations (e.g. bellbirds, *Procnias*), were given the highest score (4) because variance in the reproductive success of males is likely to reach its peak in these systems.

Unlike previous bi-directional scoring systems (Dale *et al*. 2015), we treated sexual selection as a unidirectional variable. That is, we make no distinction between male or female-biased mating systems, instead grouping polyandry and polygyny under the general definition of polygamy. To facilitate any future research that relies on discriminating between female-biased and male-biased sexual selection, we provide an updated list of sex-role-reversed bird species (Supplementary Dataset 1) so that a bidirectional score can be generated by reversing the sign on scores for sex-role-reversed species.

To enable phylogenetic analyses, we aligned sexual selection scores and associated geographical range data to the most comprehensive phylogeny available (Jetz *et al*. 2012). The taxonomy used in this phylogeny requires updating, so we also provide an additional dataset of sexual selection scores aligned with 10671 species included for a more recent taxonomy (Clements *et al*. 2021). Although we do not use this dataset in our own analyses, we hope that it will enable future studies using forthcoming phylogenetic datasets, as well as several rich sources of information, including AVONET (Tobias *et al*. 2022), Birds of the World (Billerman *et al*. 2022) and eBird citizen-science data (Sullivan *et al*. 2009), all of which use the same updated taxonomic format (Clements *et al*. 2021). Sexual selection scores under both taxonomic treatments, along with the sources of information supporting these assignments, are presented in Supplementary Dataset 1.

### Testing sexual selection metrics

To validate the accuracy of our proposed scoring system in estimating sexual selection, we conducted a comparative analysis against three widely used measures of sexual selection from existing published datasets: residual testes mass (Pitcher *et al*. 2005; Baker *et al*. 2020), Bateman gradients (β_SS_) (Fromonteil *et al*. 2023), and the opportunity for sexual selection (*I_S_*) (Janicke & Morrow 2018). Given the scarcity of studies estimating sexual selection, we also derived additional estimates of zero *I_S_* from studies that recorded 0% extrapair paternity (EPP) rates, as summarized by Brouwer & Griffith (2019). We assessed the strength of association between our scores and each metric using Bayesian phylogenetic models (Supplementary Information), with each metric as a single predictor. To test the accuracy of each metric, we extracted standard p-values for the strength of association, and calculated marginal *R*^2^ values to determine how closely scores and metrics aligned. All three metrics had significant relationships with our sexual selection scores, suggesting our scoring criteria was able to capture both pre- and post-copulatory sexual selection. To confirm a significant relationship with *I_s_* was due to estimates from species with 0% EPP, we performed a sensitivity analysis excluding the 51 species with estimated *I_s_* scores, which produced results consistent with the full models, as presented in Table S1.

### Accounting for biases in data certainty

Assembling data on sexual selection is challenging because information for many species is sparse or based on few observations. Thus, even though much information is available in standardized format (e.g. Billerman *et al*. 2022), the quality and depth of information varies greatly across species. To address these biases, we scored data certainty using a system adapted from Tobias *et al*. (2016), from 1 (lowest certainty) to 4 (highest certainty; see Extended Data Table 2 for full definitions). Species lacking direct observations receive higher certainty scores if they belong to taxonomic groups with consistent and well-documented behaviour (e.g. a poorly known species of sandgrouse would be scored 3 because all known Pteroclidae are monogamous with biparental care). Species in groups with some variation in breeding systems would instead be scored 2 because the degree of confidence in inferred data is reduced.

### Ecological variables

To assess the role of evolutionary drivers, we extracted data on species-specific traits from published datasets (Tobias *et al*. 2016, 2022). We selected three fundamental ecological traits previously proposed to influence levels of sexual selection: trophic level (linked to both spacing and constraints on parental care; Snow 1971; Beehler 1983, 1987), migration (linked to breeding synchrony or constraints on the duration of breeding seasons; García-Peña *et al*. 2009), and territoriality (linked to movement and spacing patterns; Davies & Lundberg 1984).

To facilitate the comparison of categorical and continuous data, we converted trophic level, migration, and territoriality to binary variables (Gelman 2008). For trophic level, we group species as either obligate primary consumers, defined as those with more than 60% of their energetic requirements provided by plant material, or secondary consumers, including omnivores, carnivores, and scavengers from higher trophic levels. This was because primary consumers generally experience a more heterogeneous landscape than higher trophic levels, often with scarce resources of high nutritional value. Furthermore, previous research has shown a potential relationship between herbivory and polygamy in mammals (Lukas & Clutton-Brock 2013) that may also be present within birds.

Migration was scored in three categories: sedentary, partially migratory (minority of population migrates long distance or most individuals migrate short distances), and migratory (majority of population undertakes long-distance migration). We dichotomized migration by grouping sedentary and partial migrants together, because obligate long-distance migrants tend to maintain consistent departure times, distances, and directions (Newton 2012), which should all lead to higher breeding synchronicity. Territorial behaviour was classified as either: ‘none’ (never territorial or at most defending very small areas around nest sites), ‘weak’ (weak or seasonal territoriality, including species with broadly overlapping home ranges or habitually joining mixed species flocks), and ‘strong’ (territories maintained throughout year). We dichotomized territoriality by grouping weak and strong territorial behaviour together, as both potentially create strong spatial constraints that in turn may influence the mating opportunities (Davies & Lundberg 1984; Cramer *et al*. 2020).

### Climatic seasonality

To account for the effects of climate seasonality in models of sexual selection, we included species-specific data on average temperature seasonality, extracted from the CHELSA bioclim dataset v2.1 (Karger *et al*. 2017). We selected temperature seasonality (bio4) as our primary measure of climate variability, based on extensive research connecting temperature, mating systems, and sexual selection (García-Roa *et al*. 2020; Leith *et al*. 2022). The temperature seasonality metric in CHELSA (bio4) provides an index of local intra-annual temperature variation, with grid cells equal to the standard deviation of monthly mean temperatures across each year, using global data from 1981–2010 at 30 arcsecond resolution. To extract an average seasonality estimate for each species, we overlaid climate data with expert-drawn breeding ranges provided by BirdLife International (2021) (Supplementary Information). Grid cells that fell within each species’ breeding range were averaged using a Behrmann equal area projection, disregarding cells with less than 50% overlap. Removing species without range maps, this left 9836 species for subsequent statistical analysis.

## Statistical analysis

To assess macroevolutionary patterns in sexual selection, we used Bayesian regression models from the R package *brms* (Bürkner 2017) (Supplementary Information). Given the ordinal nature of our sexual selection scores, we used a cumulative family distribution, which can be interpreted similarly to standard generalized linear models (McCullagh 1980).

Initially, we ran models including all species (*n* = 9836), with absolute latitude (distance from the equator) as the sole predictor. We also ran similar models with data certainty score as the response variable to identify any latitudinal bias in data certainty. Given a strong relationship between latitude and data certainty, we ran subsequent latitudinal models predicting sexual selection, limiting data certainty to species with moderate to high certainty (scored 3–4; *n* = 7592 species), and high certainty (scored 4; *n* = 2851 species). These models produced similar results (Extended Data Fig. 3), suggesting that our results are robust to data quality.

To determine how species ecology shapes global trends, we ran subsequent latitudinal models under the following partitions: primary consumers (*n* = 2753), frugivores (*n* = 1025), secondary consumers (*n* = 7083), invertivores (*n* = 4694), long-distance migrants (*n* = 901), resident and short-distance migrants (*n* = 8935), territorial species (*n* = 7261), and non-territorial species (*n* = 2575). To reduce the effect of spatial autocorrelation, we repeated all species-level models dichotomizing latitude as either tropical (centroid latitude within the Tropics of Capricorn and Cancer) or non-tropical. We also repeated each set of models on a high certainty subset (certainty score 3–4), which produced near-identical results (Table S3).

After identifying raw latitudinal gradients in sexual selection, we used univariate and multivariate Bayesian phylogenetic models to link potential evolutionary drivers with the strength of sexual selection across species. We first tested the phylogenetic signal for binary traits (*D*) (Fritz & Purvis 2010), dichotomizing sexual selection by grouping scores as either low sexual selection (0–2) or high (3–4). Results show that phylogenetic *D* was indistinguishable from zero (*D* = -0.19, *p* = 0.98), suggesting that sexual selection strength is highly conserved in birds and has likely evolved under a Brownian motion model. Consequently, we included a phylogenetic co-variance matrix as a random effect, using the Jetz et al. (2012) tree topology grafted to the Prum *et al*. (2015) genomic backbone. To ensure our results are robust to uncertainty in tree topology, we repeated each model over 50 randomly selected trees, combining draws into a single posterior distribution.

To determine the role of species ecology in directly predicting sexual selection, we ran phylogenetic models incorporating information on trophic level, migration, territoriality, and seasonality. As the only continuous variable, seasonality was log-transformed to approximate normality, and standardized to two standard deviations to facilitate comparison with categorical predictors (Gelman 2008). All categorical predictors were centred to reduce collinearity with interaction terms, and set the references to global averages (Schielzeth 2010). To address the hypothesis that spatial and temporal heterogeneity of resource abundance drive sexual selection, we also included tropic level interactions with seasonality (temporal heterogeneity) and territoriality (spatial heterogeneity). All predictors and interactions had VIF values below 3 (Table S5), indicating that collinearity would not affect model interpretation. To account for biases in data quality, we re-ran univariate and multivariate analyses on high quality data (certainty 3 or 4; n = 7592). All analyses were conducted in R version 4.2.1 (R Core Team 2022).

## Acknowledgements

We thank Jane Amirthanayagam, Jessamine Badcock-Scruton, Celina Chien, Anita Kristiansen, Xiaoya Lian and Thomas Munro for their help in data collection and management. Emily DuVal contributed valuable unpublished data. Illustrations from Birds of the World are reproduced with permission of Cornell Lab of Ornithology. This study was funded by a Natural Environment Research Council PhD scholarship to RAB (Doctoral Training Centre: NE/P012345/1).

## Author contributions

RAB and JAT conceived the study and managed data collection. JY, CY, OB, JD and TJ contributed data and JY helped merge datasets. RAB conducted analyses, produced figures, and wrote the first version of the manuscript, with input from JAT. All authors contributed critically to subsequent drafts and gave final permission for publication.

## Data availability

Data for all analyses are available in Supplementary Data 1. In addition, phylogenetic trees were downloaded from http://www.birdtree.org, with the specific subset of trees used in the analyses presented in Supplementary Data 2. Climate data was extracted from the CHELSA bioclim dataset v2.1 (https://chelsa-climate.org/bioclim/) and species range maps were provided by BirdLife International (http://www.datazone.birdlife.org). Silhouettes used in figures are public domain (sources given in Supplementary Data 2).

## Code availability

The R code used to run analyses and prepare figures is available at https://github.com/Syrph/sexual_selection.

## Extended data

**Extended Data Table 1.** Criteria for scoring the intensity of sexual selection in birds. All species were scored from literature by applying this classification system to textual descriptions of mating systems and breeding behaviour, including territoriality and offspring provisioning roles. By our definition, monogamy involves both sexes maintaining the same breeding partner for the duration of a reproductive event or season, and thus includes both sequential and perennial (lifelong) monogamy. Where available, we used quantitative estimates of polygamy, EPP and OSR to refine our scores. The quantitative thresholds used for scores 0–3 were based on previous studies (Verner & Willson 1966; Møller 1986; Owens & Hartley 1998). Scores for all species are presented in Supplementary Dataset 1 along with literature sources and estimates of data certainty (see Extended Data Table 2).

| Score | Criteria |
| --- | --- |
| 0 | Strict monogamy, including extreme low rates (<0.1%) of individual polygamy and <5% extra-pair paternity (EPP). |
| 1 | Frequent monogamy, including relatively low rates (0.1–5%) of individual polygamy and <25% EPP. Species with occasional or anecdotal records of polygamy were assigned this score. |
| 2 | Regular polygamy, including cases with multiple records of polygamy. Where empirical data are available, this score was applied to species with 5–20% of individuals polygamous. This score was also applied to species with 25–50% EPP, or highly skew in operational sex ratio (OSR). |
| 3 | Frequent polygamy with >20% of individuals polygamous or very high rates of EPP (>50% of offspring) or extremely skewed OSR. |
| 4 | Extreme polygamy characterized by communal display (lekking) behaviour or permanent display posts used by solitary individuals. This score was not assigned according to levels of polygamy, EPP or OSR. |

**Extended Data Table 2.** Scoring system for assigning data certainty to estimates of sexual selection in birds. Bird species vary from well-known to almost entirely unknown in life, creating potential biases in knowledge. To account for this, we applied a classification system to quantify the availability of information relevant to the scoring procedure for estimating levels of sexual selection in each species. This classification system is adapted from Tobias *et al*. (2016).

| Score | Criteria |
| --- | --- |
| 1 | No direct or indirect evidence, including a lack of clear information preventing inference from closely related species. This score is mainly restricted to extremely rare species with no close relative, or poorly known species occurring in taxonomic groups with no clear pattern in mating systems. |
| 2 | Low levels of certainty due to conflicting evidence. This includes cases where inference from closely related species is complicated by high levels of variation in mating systems or where published evidence about mating systems is conflicting. |
| 3 | Moderate levels of certainty based on inference and reasonable circumstantial evidence. This includes reliable field observations suggestive of particular mating systems, as well as inference from closely related species. |
| 4 | High levels of certainty based on direct evidence published in primary and secondary literature. |

**Extended Data Fig. 1.**
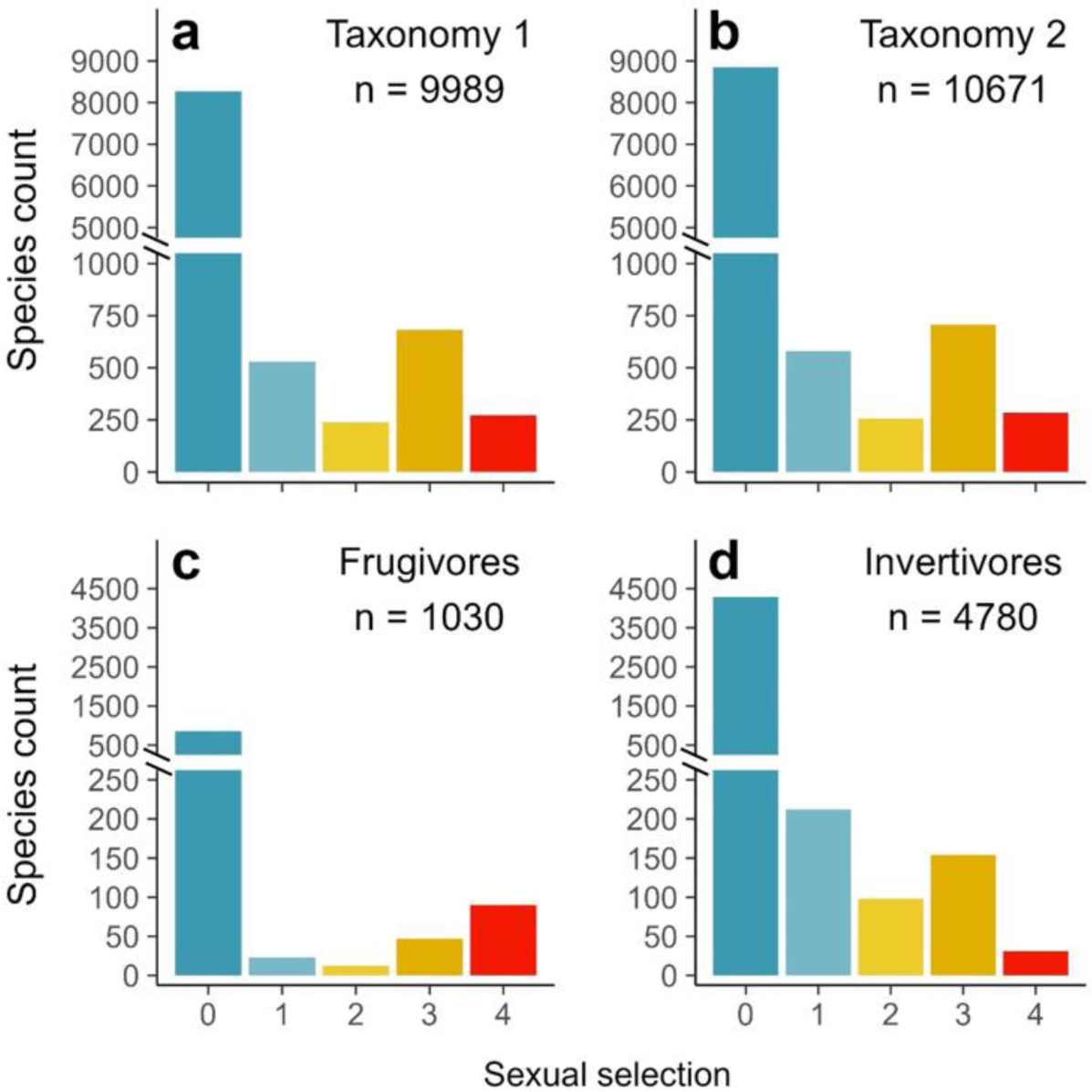
Species richness of birds assigned to different levels of sexual selection. Plots show number of species in categories 0–4 according to the species limits in BirdTree (Jetz *et al*. 2012) (**a**) or a more recent taxonomic update (Clements *et al*. 2021) (**b**), as well as within the two largest dietary guilds: invertivory (**c**) and frugivory (**d**). The strength of sexual selection increases from 0 (strict monogamy) to 4 (extreme polygamy) according to our scoring system (Extended Data Table 1).

**Extended Data Fig. 2.**
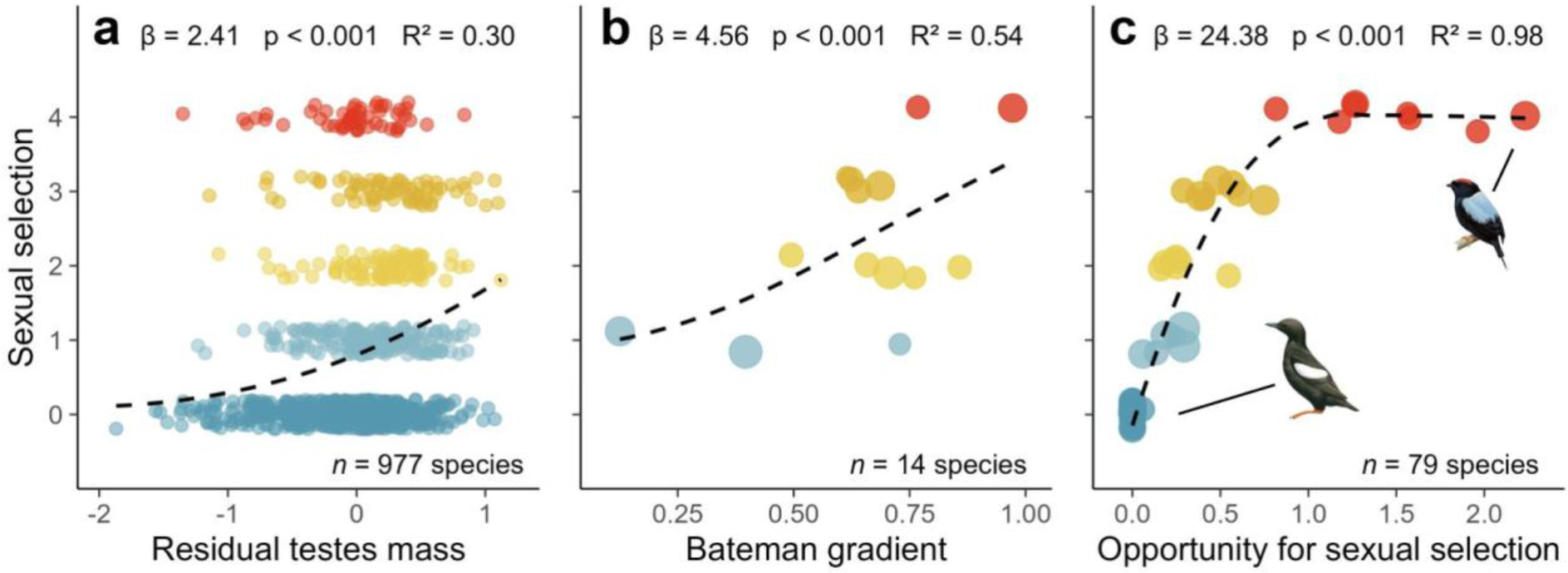
Comparison of scores with alternative metrics of sexual selection. Panels show results of Bayesian phylogenetic models assessing the relationship between our sexual selection scores and three independent measures of sexual selection: (**a**) residual testes mass, (**b**) Bateman gradients (β_ss_), (**c**) the opportunity for sexual selection (*I_s_*) (see Methods). Dashed lines show the relationship for subsets of species with available data, using predictions from models accounting for phylogeny using a sample of 50 phylogenetic trees extracted from www.birdtree.org (Jetz *et al*. 2012), grafted to the Prum *et al*. (2015) genomic backbone. Effect sizes, p-values, and explained variance (*R*^2^) were calculated from fixed effects only, and therefore represent the specific variation explained by each predictor separate from phylogenetic effects.. In **b** and **c**, point size is scaled by the number of individuals used to calculate sex-specific metrics. To compare sex-specific metrics with bidirectional scores, we selected the largest β_ss_ or *I_s_* from each species, treating males and female metrics equally. We estimated *I_s_* of 51 genetically monogamous species (with 0% extra-pair paternity) as zero. Excluding these species from *I_s_* models produced similar results (Table S1). Illustrations of Black Guillemot and Lance-tailed Manakin are shown as exemplars of extremely low and extremely high *I_s_*, respectively; images shared with permission of Birds of the World, Cornell University.

**Extended Data Fig. 3.**
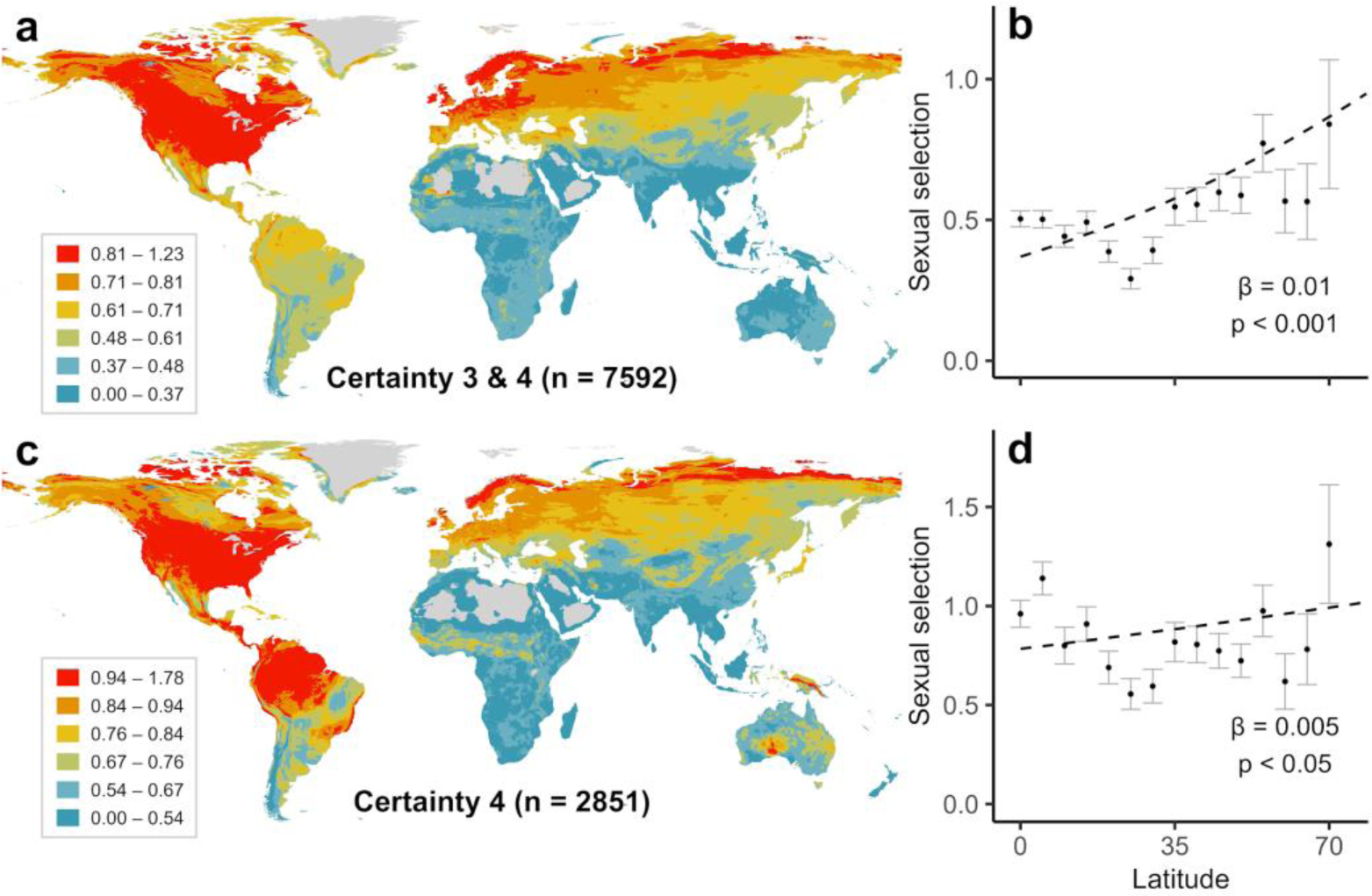
Effects of data quality on the distribution of sexual selection. **a**, Upper panels show average sexual selection for species with moderate to high data certainty (scored 3–4; *n* = 7592) mapped globally (**a**) and plotted against latitude (**b**). Lower panels show average sexual selection for species with high certainty data only (scored 4; *n* = 2851), again mapped globally (**c**) and plotted against latitude (**d**). In **a** and **c**, averages for each cell are calculated from all species with geographical range maps overlapping each 5-km grid cell; cells with <10 species were excluded. In **b** and **d**, sexual selection scores are averaged across five-degree latitudinal bins, with each species assigned to a single bin according to its latitudinal centroid; and latitudinal bins above 70 degrees were excluded from all scatterplots. Points represent means and range bars denote one standard error; dashed lines were generated from species-level Bayesian regression models predicting sexual selection strength.

**Extended Data Fig. 4.**
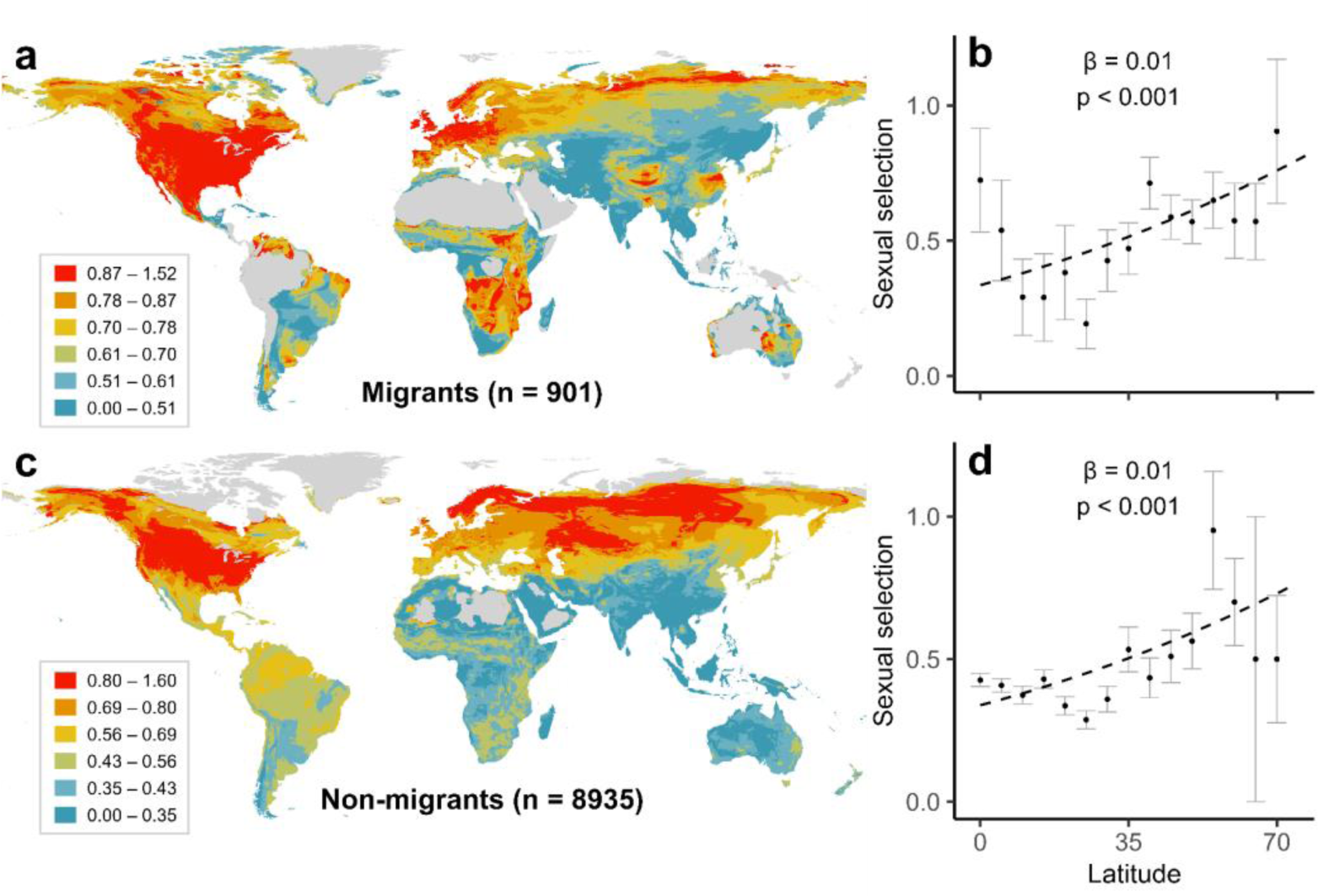
Global distribution of sexual selection partitioned by migration tendency. Upper panels show strength of sexual selection in long-distance migrants mapped globally (**a**) and plotted against latitude (**b**). Lower panels show strength of sexual selection in short-distance and resident species mapped globally (**c**) and plotted against latitude (**d**). Sexual selection was scored in five categories ranging from monogamy (0) to extreme polygamy (4; see Methods). In **a** and **c**, averages for each cell are calculated from all species with geographical range maps overlapping each 5-km grid cell; In scatterplots, variables were averaged within five-degree latitudinal bins based on the centroid latitude of their geographical ranges; points represent means; bars denote one standard error; dashed lines were generated from species-level Bayesian regression models predicting sexual selection strength. Additional species-level models on a high certainty subset (scored 3–4) showed similar patterns and are reported in Table S2. To reduce noise, cells with <10 species were excluded from all maps, and latitudinal bins above 70 degrees were excluded from all scatterplots.

**Extended Data Fig. 5.**
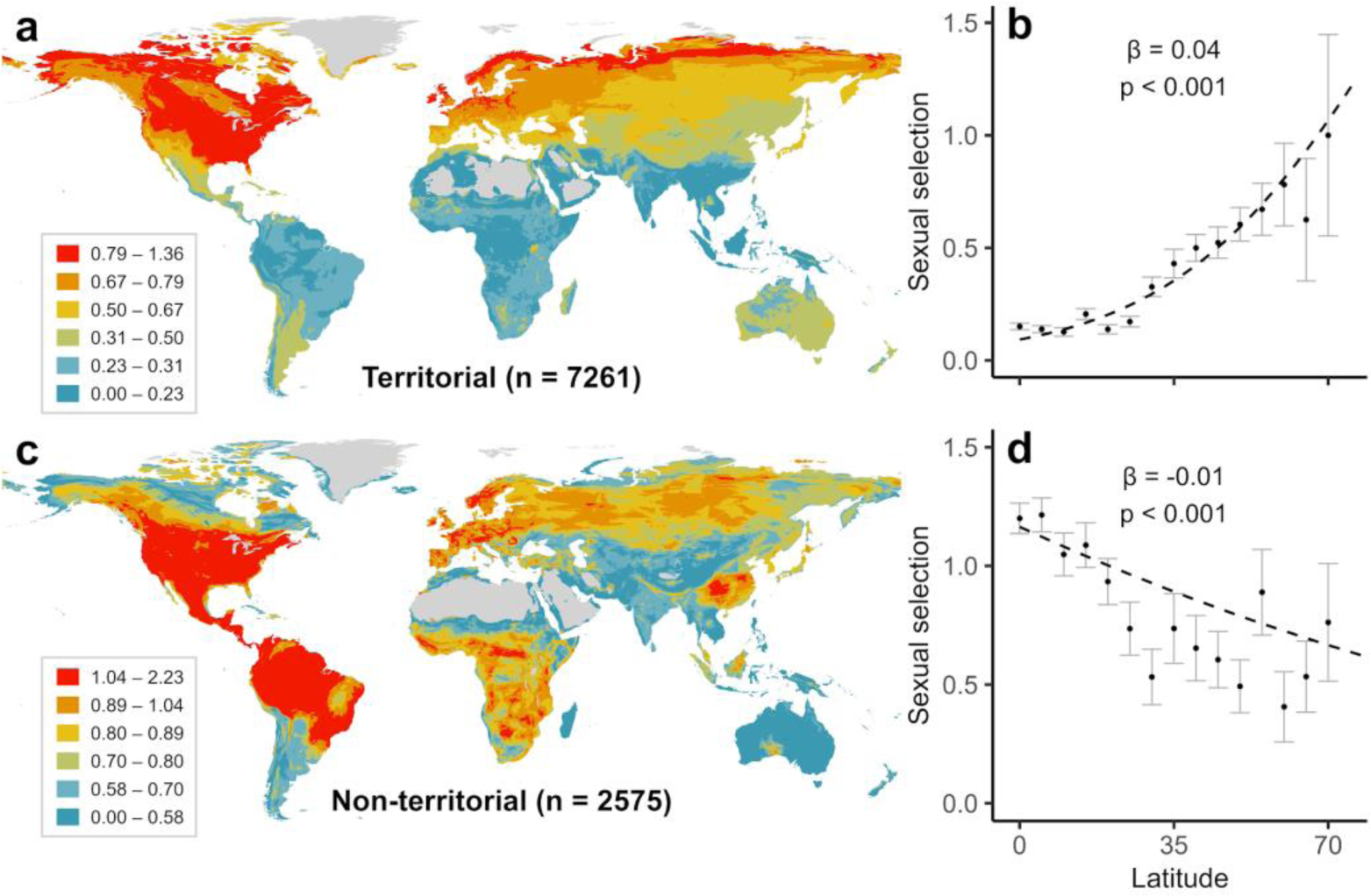
Global distribution of sexual selection partitioned by territorial behaviour. Upper panels show strength of sexual selection in seasonal and year-round territorial species mapped globally (**a**) and plotted against latitude (**b**). Lower panels show strength of sexual selection in non-territorial species mapped globally (**c**) and plotted against latitude (**d**). Sexual selection was scored in five categories ranging from monogamy (0) to extreme polygamy (4; see Methods). In **a** and **c**, averages for each cell are calculated from all species with geographical range maps overlapping each 5-km grid cell. In scatterplots, variables were averaged within five degree latitudinal bins based on the centroid latitude of their geographical ranges; points represent means; bars denote one standard error; dashed lines were generated from species-level bayesian regression models predicting sexual selection strength. Additional species-level models on a high certainty subset (scored 3–4) showed similar patterns and are reported in Table S2. To reduce noise, cells with <10 species were excluded from all maps, and latitudinal bins above 70 degrees were excluded from all scatterplots.

**Extended Data Fig. 6.**
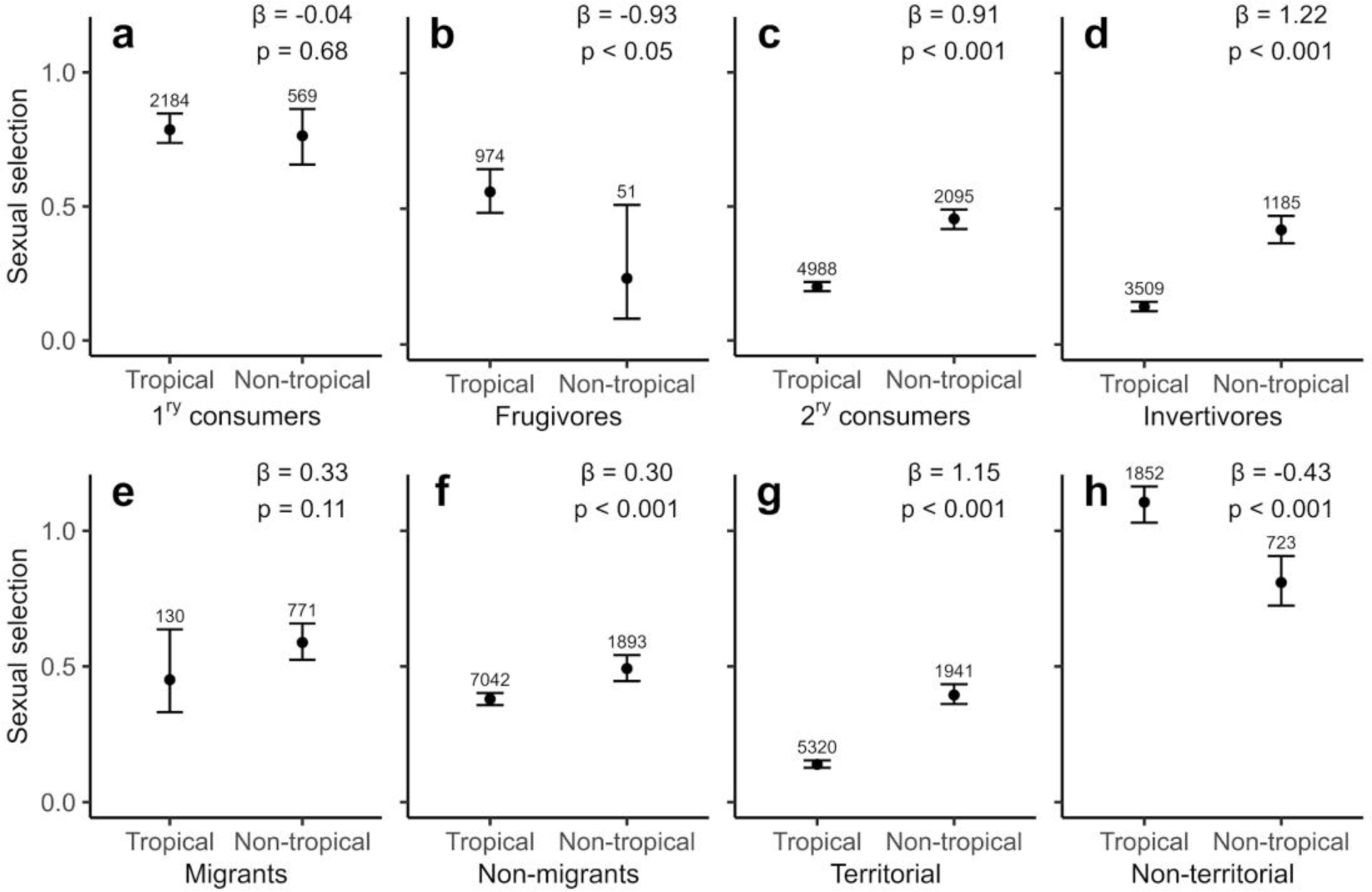
Variation in sexual selection for tropical and non-tropical species. Models predicting the difference in sexual selection strength between tropical and non-tropical species, partitioned by ecological trait. Black points show estimated sexual selection for each group, extracted from species-level bayesian regression models; whiskers show 95% confidence intervals from model predictions. Additional species-level models on a high certainty subset (scored 3–4) showed similar patterns (Table S3; Supplementary Information). Numbers above whiskers are sample sizes (number of species).

**Extended Data Fig. 7.**
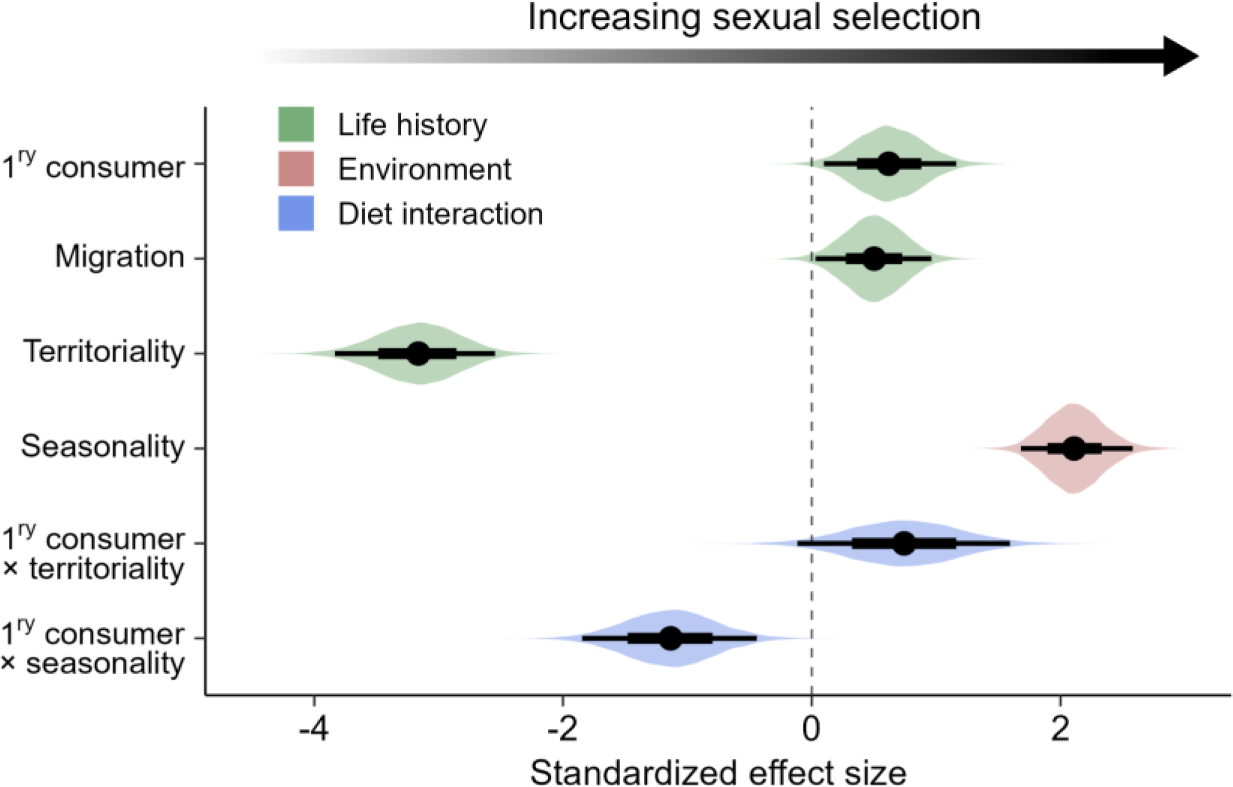
Evolutionary drivers of avian sexual selection. Results shown are from Bayesian phylogenetic models testing drivers of sexual selection in species scored with moderate and high data certainty scores (scored 3–4; *n* = 7592 species). Predictors include three life history variables (green), one climatic variable (pink), and two key interactions between diet and the dominant effects (territoriality and seasonality; blue). The reference groups for the three categorical predictors are as follows: secondary consumer; no migration; and no territoriality, respectively (see Methods for definitions). Models were run on a sample of 50 phylogenetic trees extracted from www.birdtree.org (Jetz *et al*. 2012), grafted to the Prum *et al*. (2015) genomic backbone. Dots show mean effect size estimates from 12,500 posterior draws. For each effect, broad bases of whiskers show 66% confidence intervals; narrow tips of whiskers show 95% confidence intervals. Coloured distributions indicate the spread of effect size estimates, generated from a sample of 1000 posterior draws. Full statistical results are presented in Supplementary Table S5.

**Extended Data Fig. 8.**
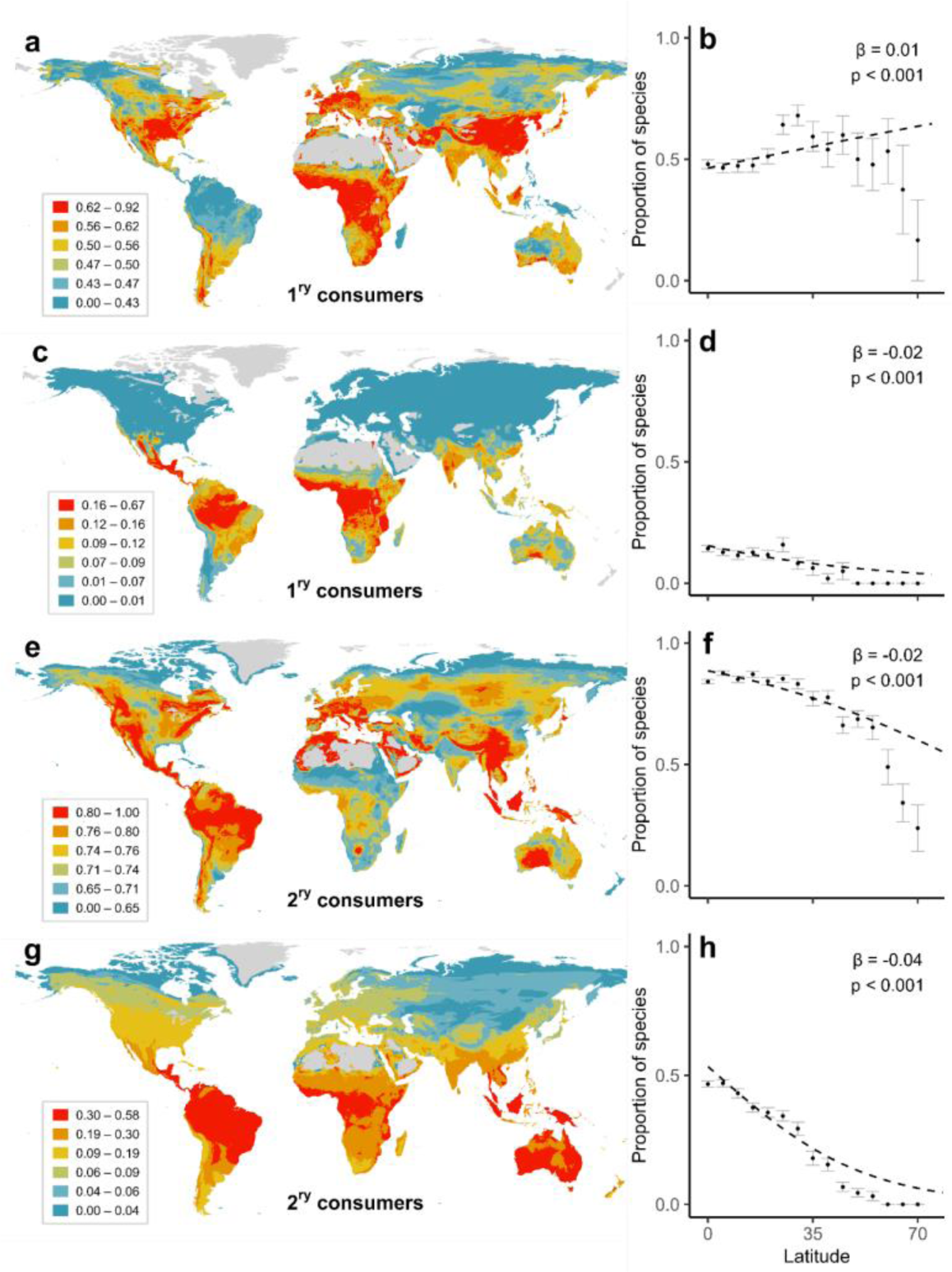
Biogeography of avian territoriality. Proportion of territorial primary consumers (*n* = 1379 species) mapped globally (**a**), and plotted against latitude (**b**), compared with those holding year-round territories only (**c**, **d**; *n* = 333 species). Proportion of territorial secondary consumers (*n* = 5882 species) mapped globally (**e**) and plotted against latitude (**f**), compared with those holding year-round territories only (**g**, **h**; *n* = 2640 species). Territoriality data (see Methods) were converted into binary scores for mapping purposes, as follows: in **a, b, e** and **f** (0 = none; 1 = seasonal/year-round); in **c, d, g** and **h** (0 = none/seasonal; 1 = year-round). In **a**, **c**, **e** and **g** averages for each 5-km grid cell are calculated from all species with geographical range maps overlapping each cell. In **b**, **d**, **f** and **h**, scores were averaged in 5-degree bins based on the centroid latitude of ranges; latitudinal bins above 70 degrees were excluded from all scatterplots. Points represent means; bars denote one standard error; dashed lines were generated from species-level Bayesian regression models. In **a**, **c**, **e** and **g**, cells with <10 species were excluded from maps.

### SUPPLEMENTARY INFORMATION

As separate Excel file: Supplementary dataset 1

Sheet 1 – Metadata. Definitions and sources for all variables in subsequent data sheets.

Sheet 2 – Data1 (BirdTree). Sexual selection scores and associated uncertainty estimates and source citations for 9988 bird species included in the BirdTree dataset (Jetz *et al*. 2012), along with corresponding taxonomic, ecological, morphological and distributional data, where available (*n* = 9863 species).

Sheet 3 – Data2 (Clements). Sexual selection scores aligned with an updated taxonomy (Clements *et al*. 2021) used in major online resources, including Birds of the World and eBird (*n* = 10671 species).

Sheet 4 - Sexual selection metrics, detailing residual testes mass, bateman gradients, and opportunity for sexual selection values used in sensitivity analysis.

Sheet 5 – Taxonomic crosswalk. Guide to species matching for users transferring data between BirdTree and Clements (eBird) taxonomies, which often differ in spelling and treatment of allospecies.

Sheet 6 – Data sources. Full references for sources cited in Data1 (BirdTree).

### Supplementary methods

#### Taxonomy

To allow the use of phylogenetic analyses, we followed the species names used in the global BirdTree phylogeny (Jetz *et al*. 2012). Given the rapid state of flux in avian taxonomy, the species limits and family classifications in BirdTree are increasingly obsolete, particularly because the phylogenetic tree itself is likely to be replaced with new phylogenies using updated taxonomic treatments. To maximise future utility of our dataset, we rescored sexual selection and all other variables according to Clements (2021) taxonomy. This will allow integration of our dataset with ecological and life history data published in Birds of the World (https://birdsoftheworld.org/bow), as well as spatially explicit citizen-science data from eBird (https://ebird.org), and forthcoming global phylogenetic datasets based on Clements taxonomy.

Approximately 93% of species in the BirdTree dataset were directly matched with Clements species, including many that needed to be matched manually because of changes in spelling or higher taxonomic classification. In all these cases, we transferred the same sexual selection and data uncertainty scores from BirdTree to the Clements dataset. For the remaining 7% of species, 512 were taxonomic splits, where a BirdTree species was separated into multiple Clements species. In these cases, we identified the parent (nominate) species and transferred trait data directly from BirdTree to the parent lineage. For the remaining daughter species, we inferred sexual selection scores from the behaviour of the parent. For example, under BirdTree taxonomy, the House Sparrow *Passer domesticus* is considered a single species, with the Italian Sparrow considered a subspecies (*P. domesticus italiae*). However, under Clements taxonomy, *P. italiae* is considered a separate species. In this instance, we directly transferred sexual selection and data certainty scores from the BirdTree “*P. domesticus”* values to the Clements “*P. domesticus*” values. For *P. italiae*, we used the same sexual selection score as the parent by inference, because breeding behaviour of this geographically restricted form appears similar to the more widespread *P. domesticus*. Nonetheless, we dropped the data certainty score from 4 to 3 reflecting the reduction in certainty that arises because *P. italiae* occupies a small geographical range and the sexual selection score is no longer based on direct species-specific evidence.

The reverse procedure was required in cases where multiple BirdTree species have been merged into a single Clements species. These cases were less frequent, with only 219 BirdTree species combined into 102 Clements species. In almost all cases, sexual selection scores were the same across each set of BirdTree species, so this score was directly transferred to the corresponding Clements species into which they were merged. The only exception was the Ring-necked Pheasant (*Phasianus colchicus*), in which two BirdTree sister species were scored differently: the mainland nominate *P. colchicus* was scored 3 for polygamy, whereas the Green Pheasant (*P. versicolor*), a Japanese endemic, was scored 2 because a higher incidence of monogamy was reported than for mainland populations. Under 2021 Clements taxonomy, *P. colchicus* was merged with *P. versicolor* and took the higher score (3) because the Japanese form makes up only a small part of the total population.

When transferring data from BirdTree to Clements, 31 species presented challenges because data certainty scores differed between daughter taxa being merged together (lumped) into a single parent taxon. In most of these problematic cases, the sexual selection score of widespread parent taxa was based on direct data from primary sources, whereas this direct data often only applied to one of the BirdTree daughter species (splits). In these instances, we transferred the certainty data from the nominate BirdTree species to the lumped (parent) eBird species. The only exception to this rule was *Stercorarius antarcticus* which is considered two separate species (*Catharacta lonnbergi* and *Catharacta antarctica*) by BirdTree and a single species by Clements. Under BirdTree, the daughter species or split, *C. lonnbergi*, carries a higher data certainty score (4) because its sexual selection score is based on a molecular study of extra-pair paternity (Millar 1994). In the same treatment, *C. antarctica* has a lower certainty (3) which is upgraded to higher certainty (4) under Clements taxonomy wherein *C. lonnbergi* is subsumed into *C. antarctica* (and transferred to the genus *Stercorarius*).

In five cases, BirdTree species were initially lumped and then subsequently re-split into multiple daughter species. However, sexual selection happened to be scored as strict monogamy (0) in all these cases, simplifying the direct transfer of scores from BirdTree to Clements datasets. Finally, 176 Clements species had no corresponding BirdTree species. The majority of cases (*n* = 144 species) were extinct species that were omitted from BirdTree, which we did not include in our database. The remaining 32 species were described in the last decade, representing new additions to global species lists. In these cases, we scored sexual selection from literature following the same protocol used for BirdTree species (see Methods).

We include a taxonomic crosswalk (sheet 4) in Supplementary dataset 1. This resource lists each BirdTree and Clements species aligned in neighbouring columns, showing how they are matched in our data transfer process. Specifically, we include the type of link between the two taxonomies (one-to-one, many-to-one, one-to-many, many-to-many) and highlight the parent taxon in cases where data are duplicated across multiple taxa in the case of splits. The crosswalk is designed to be used alongside the R script (clements_conversion.R) included with this study. As a disclaimer, our crosswalk may contain small errors as it was designed to transfer traits between thousands of species. We recommend that users carefully check the conversion of traits, paying particular attention to species which have transitioned between families or orders. We hope that the Clements dataset and associated taxonomic crosswalk can support future researchers by facilitating the transfer of traits to and from Clements taxonomy, including cases where lumped BirdTree species have conflicting data. This should be a useful resource for macroecological studies combining data from AVONET (Tobias *et al*. 2022), Birds of the World, and eBird, or for researchers wishing to integrate their datasets with future phylogenies based on Clements taxonomy.

#### Sexual selection metrics

To assess the reliability of our scoring system, we collected data on three complementary and widely used measures of sexual selection from existing published datasets: residual testes mass (Pitcher *et al*. 2005; Baker *et al*. 2020), Bateman gradients (β_SS_) (Janicke *et al*. 2016; Fromonteil *et al*. 2023), and the opportunity for sexual selection (*I_S_*) (Janicke & Morrow 2018). Residual testes size constitutes a commonly used proxy for the strength of post-copulatory sexual selection on males whereas β_SS_ and *I_S_* are standardized metrics for the strength of pre-copulatory sexual selection. To increase our sample size, we also calculated metrics for additional species where possible, based on published data.

Residual testes mass refers to the mass of testes after accounting for total body mass. This metric provides information about the levels of sperm competition between rival males, which correlates with some aspects of sexual selection (Lüpold *et al*. 2020). To calculate residual testes mass, we used recent data from Baker *et al*. (2020) (n = 901 species), with an additional 76 species sourced from Pitcher *et al*. (2005). Because Baker *et al*. (2020) published testes mass and body mass as separate measurements, we calculated residual testes mass under the same procedure as Pitcher *et al*. (2005), using the residuals from the regression of log testes mass on log body mass.

*I_S_* can be calculated as the sex-specific variance in relativized mating success across individuals of a population (defined as the variance in the number of sexual partners per individual, divided by the squared mean of the number of sexual partners). We calculated *I_S_* for 28 unique species, either directly from information presented in papers, or from data subsequently provided by authors. Given the scarcity of studies that calculate quantitative measures of sexual selection, we also estimated *I_S_* for 51 monogamous species that recorded 0% extrapair paternity (EPP), indicating true genetic monogamy. Assuming that each individual was paired, and that operational sex-ratios are approximately even (as tends to be true for genetically monogamous species), this meant we could assign an *I_S_* of zero for these 51 additional species. As a sensitivity analysis, we removed these species from our reliability assessment (see main Methods), which produced near-identical results (Table S1).

Bateman gradients (also termed sexual selection gradients) represent the slope of an ordinary least squares regression of reproductive success (number of offspring) on mating success (typically quantified as the number of mating partners) and therefore estimate the fitness benefit that can be obtained from additional matings. To compare between species, we converted Bateman gradients into standardised Pearson correlation coefficients. Lastly, because our sexual selection scores are bi-directional (scoring polyandry and polygyny equally), we took the recorded *I_S_* and β_SS_ from the sex with the highest value to generate a single measure per species.

#### Spatial methods

To understand how sexual selection varies globally, we incorporated spatial information using expert-drawn geographical range maps provided by Birdlife International (2021). To align species with BirdTree (Jetz *et al*. 2012), we used the BirdLife-to-BirdTree taxonomic crosswalk provided in AVONET (Tobias *et al*. 2022). In most cases, BirdTree species had a single equivalent BirdLife species name (one-to-one matches; *n* = 8949), which allowed us to use the corresponding geographical range polygon for the same species.

The majority of taxonomic differences between BirdTree and BirdLife were splits, which resulted in a single BirdTree species corresponding with multiple BirdLife species (BirdLife splits; *n* = 1929). In these cases, we combined multiple BirdLife daughter species into a single range polygon for their corresponding parent BirdTree species using the *sf* package (Pebesma 2018). In the small number of cases where multiple BirdTree species correspond with one BirdLife species (BirdLife lumps; *n* = 198), we used BirdLife version 2.0 maps (BirdLife International 2012), created prior to any taxonomic changes. In this resource, we used maps for the corresponding BirdLife synonyms for each BirdTree species.

For a small number of BirdLife lumps (*n* = 64), no previous geographical range polygon existed as a version 2.0 map. In addition, some taxonomically stable species lacked geographical data (*n* = 57), either because no part of their known range is coded as resident or breeding range, or because the map is not openly released by BirdLife International (typically because the species is sensitive to hunting or trapping for the cage-bird trade). For mapping purposes, we restricted range polygons to areas where the species is coded by BirdLife International as extant and either native or reintroduced. Finally, because latitude and environmental conditions in the non-breeding range are largely irrelevant to levels of sexual selection, we limited our selection to the breeding and resident ranges of each study species.

To extract species-specific data, we used a Behrmann equal-area projection for geographical range polygons. To calculate a single value of latitude for each species, we determined the geometric centroid of each range polygon using the *PBSmapping* package (Schnute *et al*. 2017). This approach was preferred over midpoint latitude metrics because the centroid considers the overall shape of a species’ distribution, particularly important for wide-ranging species. Assigning a single value of latitude to each species greatly over-simplifies their distribution, and can be misleading, particularly in species with wide ranges spanning the equator. Nonetheless, midpoint or centroid estimates have the advantage of simplicity, removing the statistical problem of repeated measures per taxon, and usually provide a reasonable approximation of the latitude at which most individuals in the global population breed. For these reasons, midpoints or centroids are often used in phylogenetic comparative analyses or macroevolutionary models testing for latitudinal effects (Weir & Schluter 2007; Sheard *et al*. 2020; Weeks *et al*. 2022).

To visualise global patterns of sexual selection, we first created a unified map by combining species ranges into a single raster by overlapping the range with a grid of cells at 5-km resolution. To calculate average values of sexual selection per cell, we summed the sexual selection scores of all species with geographical ranges overlapping more than 50% of that cell, then divided the total by the corresponding species richness of the cell. This produced a single raster with cell values representing average sexual selection for the species present. We repeated this process using data certainty scores to generate a map of average data certainty for all birds and for each ecological partition listed in Table S2.

#### Bayesian ordinal models

Bayesian mixed-effect models were constructed using the brms package in R (Bürkner 2017). Because our sexual selection scores are rankings, we used ordinal regression models with a cumulative family distribution. This model estimates the cumulative odds of being at or below each category level, providing insights into how the independent variables influence the likelihood of moving up or down the ordinal scale. Coefficient estimates and confidence intervals can be interpreted similarly to other standard generalised-linear models. Posterior predictive checks (Fig. S2) confirmed that a cumulative family distribution fits the data extremely well.

To first establish if there were latitudinal gradients in sexual selection, we used univariate models with sexual selection score as the response and absolute latitude as the predictor. We chose not to apply any additional transformations to latitude because models fit generally well, and this allowed for the best alignment with maps of sexual selection. We first ran a single model predicting sexual selection strength for all species (*n* = 9836), which showed a strong positive gradient in sexual selection (Table S2). Given the strong positive effect of latitude on sexual selection across species, we ran subsequent univariate models to determine if the overall pattern was true for different ecological groups, using the same response and predictor for different data partitions. We then repeated each model using a high certainty dataset (certainty score 3 – 4), which produced similar results. To reduce spatial autocorrelation, we performed a sensitivity analysis by dichotomizing latitude as either tropical (centroid latitude within the Tropics of Capricorn and Cancer) or non-tropical. We then used our binary latitudinal predictor to repeat the set of models outlined in Table S2, which produced similar results (Table S3).

For univariate and multivariate trait models, designed to identify the evolutionary determinants of sexual selection strength, we included a phylogenetic covariance matrix as a random effect, using the Jetz *et al*. (2012) tree topology grafted to the Prum *et al*. (2015) genomic backbone. Temperature seasonality – the only continuous variable – was log-transformed to approximate normality and standardised to two standard deviations to facilitate comparison with categorical predictors (Gelman 2008). All categorical predictors were centred to reduce collinearity with interaction terms, and to set the reference level for each trait to the global average (Schielzeth 2010). The variance inflation factors (VIF) of all model terms were below three, suggesting that collinearity between predictors does not affect model interpretation (Table S3). To further address issues of collinearity, multivariate models used additional QR decomposition (whereby a matrix is expressed as the product of two separate matrices, Q and R). This approach helps recover accurate coefficient estimates in the presence of correlated traits (Bürkner 2017).

For all Bayesian ordinal models, we used a no u-turn sampler (NUTS) and selected 10,000 total iterations with a 5,000-iteration warmup-phase and thinning every 20 iterations. Each model was run with two chains which we assessed for convergence by visually inspecting the mixing of chains and ensuring that r-hat values reached a value of one. To ensure our results are robust to uncertainty in tree topology, we repeated each model including phylogeny over 50 randomly selected trees, combining draws into a single posterior distribution. Following recommendations in Gelman (2006), we assigned weakly informative priors to the intercept and slope parameters: normal (0,1). To improve sampling speed, and because initial models showed the estimate for the phylogenetic effect was consistently close to 1, we assigned a stronger prior to the phylogenetic effect: gamma (2,1). To assess the proportion of variance explained by model predictors, we used approximate *R^2^* values for Bayesian methods (Gelman *et al*. 2019), adapted to fit ordinal regression models (McKelvey & Zavoina 1975). To calculate reported p-values from Bayesian regression models, we extracted the probability of direction from posterior draws, as described by Makowski *et al*. (2019).

#### Supplementary discussion

We considered using information about parental care roles as evidence for the strength of sexual selection. Costly parental care is often cited as a major hypothesis explaining monogamy (Klug 2018; Kvarnemo 2018) because time constraints can prevent individuals from seeking additional mating opportunities. We excluded parental care in our models because data is lacking at the scale required to test global hypotheses in birds. Alternative binary traits with good coverage such as developmental mode are unsuitable because large sections of the tree have no variation, limiting their accuracy in phylogenetic models.

#### Supplementary tables

**Table S1.** Comparing alternative metrics of sexual selection. Results shown are Results are β estimates and 95% credible intervals from phylogenetic Bayesian ordinal mixed-effect models estimating the strength of relationship between sexual selection scores from this study and alternative measures of sexual selection: residual testes mass (a proxy for post-copulatory sexual selection), Bateman gradients (β_SS_) (reproductive benefit from additional matings), the opportunity for sexual selection (*I_S_*) (variance in mating success). To compare sex-specific metrics with bi-directional scores, we selected the largest β_SS_ or *I*_S_ from each species, treating males and female metrics equally. For 51 monogamous species with 0% extra-pair paternity reported in well-sampled molecular studies, we scored *I_S_* as zero. When we ran a sensitivity analysis (*I_S_*2) with these arbitrary zeroes excluded, results remained similar. Models were run on a sample of 50 phylogenetic tree topologies extracted from www.birdtree.org (Jetz *et al*. 2012), grafted to the Prum *et al*. (2015) genomic backbone. Testes mass = residual testes mass. See Supplementary methods for detailed model descriptions.

| Metric | *n* | Estimate | 95% CI | Conditional $R^2$ | Marginal $R^2$ |
| --- | --- | --- | --- | --- | --- |
| Testes mass | 977 | 2.41 | 1.69, 3.36 | 0.91 | 0.30 |
| $\beta_{ss}$ | 14 | 4.51 | 0.55, 10.86 | 0.76 | 0.53 |
| $I_s$ | 79 | 24.08 | 15.48, 37.94 | 0.98 | 0.97 |
| $I_{s2}$ | 28 | 24.24 | 15.59, 38.47 | 0.98 | 0.97 |

**Table S2.** Latitudinal gradients in sexual selection. Results shown are β estimates and 95% credible intervals from Bayesian ordinal mixed-effect models estimating the strength of latitudinal gradients in sexual selection across different ecological groups of birds. Models were repeated on high certainty data, defined as species scored 3 or 4 for data certainty (see Extended Data Table 2). See Supplementary methods for detailed model descriptions. To increase legibility, estimates were multiplied by 10 to represent the change in sexual selection score over a 10 degree increase in latitude. See Figs. 2 & 3, and Extended Data Figs. 3 & 4 for visualisation (main text).

| Sample | Certainty 1-4 |  |  | Certainty 3-4 |  |  |
| --- | --- | --- | --- | --- | --- | --- |
|  | *n* | Estimate | 95% CI | *n* | Estimate | 95% CI |
| All birds | 9836 | 0.17 | 0.14, 0.20 | 7592 | 0.15 | 0.12, 0.18 |
| 1<sup>ry</sup> consumers | 2753 | 0.01 | -0.04, 0.07 | 2311 | -0.03 | -0.09, 0.03 |
| Frugivores | 1025 | -0.24 | -0.47, -0.01 | 767 | -0.30 | -0.57, -0.05 |
| 2<sup>ry</sup> consumers | 7083 | 0.30 | 0.26, 0.34 | 5281 | 0.32 | 0.28, 0.36 |
| Invertivores | 4694 | 0.45 | 0.39, 0.51 | 3477 | 0.56 | 0.50, 0.63 |
| Migrants | 901 | 0.15 | 0.06, 0.23 | 793 | 0.14 | 0.05, 0.23 |
| Non-migrants | 8935 | 0.13 | 0.09, 0.17 | 6799 | 0.11 | 0.06, 0.15 |
| Territorial | 7261 | 0.42 | 0.37, 0.46 | 5353 | 0.42 | 0.37, 0.47 |
| Non-territorial | 2575 | -0.11 | -0.15, -0.06 | 2239 | -0.12 | -0.17, -0.08 |

**Table S3.** Differences in sexual selection for tropical and non-tropical species. Results shown are β estimates and 95% credible intervals from Bayesian ordinal mixed-effect models estimating the difference in the strength of sexual selection between tropical and non-tropical species for different ecological groups of birds. Species were assigned tropical status if the centre of their range was within the Tropics of Capricorn and Cancer (< 23.4 degrees latitude). Models were repeated on high certainty data, defined as species scored 3 or 4 for data certainty (see Extended Data Table 2). See Supplementary methods for detailed model descriptions. See Extended Data Fig. 6 for visualisation (main text).

| Sample | Certainty 1-4 |  |  | Certainty 3-4 |  |  |
| --- | --- | --- | --- | --- | --- | --- |
|  | *n* | Estimate | 95% CI | *n* | Estimate | 95% CI |
| 1<sup>ry</sup> consumers | 2753 | -0.04 | -0.24, 0.14 | 2311 | -0.18 | -0.41, 0.03 |
| Frugivores | 1025 | -0.95 | -1.94, -0.16 | 767 | -1.06 | -2.04, -0.22 |
| 2<sup>ry</sup> consumers | 7083 | 0.91 | 0.77, 1.05 | 5281 | 0.99 | 0.85, 1.14 |
| Invertivores | 4694 | 1.21 | 1.04, 1.39 | 3477 | 1.52 | 1.3, 1.74 |
| Migrants | 901 | 0.33 | -0.11, 0.75 | 793 | 0.43 | -0.04, 0.9 |
| Non-migrants | 8935 | 0.30 | 0.17, 0.42 | 6799 | 0.21 | 0.07, 0.34 |
| Territorial | 7261 | 1.15 | 1.00, 1.30 | 5353 | 1.19 | 1.01, 1.35 |
| Non-territorial | 2575 | -0.43 | -0.61, -0.23 | 2239 | -0.52 | -0.69, -0.33 |

**Table S4.** Ecological predictors of sexual selection. Results shown are β estimates and 95% credible intervals from univariate and multivariate phylogenetic Bayesian ordinal mixed-effect models testing drivers of sexual selection in birds (*n* = 9836 species). Models were run on a sample of 50 phylogenetic trees extracted from www.birdtree.org (Jetz *et al*. 2012), grafted to the Prum *et al*. (2015) genomic backbone. See Supplementary methods for detailed model descriptions. See Figs. 4 & 5 (main text) for visualisation. 1^ry^ consumer = primary consumer.

| Predictor | Univariate |  | Multivariate |  |
| --- | --- | --- | --- | --- |
|  | Estimate | 95% CI | Estimate | 95% CI |
| 1<sup>ry</sup> consumer | 0.52 | 0.13, 0.92 | 0.64 | 0.21, 1.07 |
| Migration | 2.06 | 1.70, 2.43 | 0.84 | 0.44, 1.25 |
| Territoriality | -2.61 | -3.13, -2.12 | -2.71 | -3.27, -2.17 |
| Seasonality | 2.12 | 1.83, 2.43 | 1.78 | 1.44, 2.12 |
| 1<sup>ry</sup> consumer × territoriality | - | - | 0.80 | 0.08, 1.51 |
| 1<sup>ry</sup> consumer × seasonality | - | - | -0.75 | -1.33, -0.18 |

**Table S5.** Results of sensitivity analyses assessing robustness of models to data certainty. Results shown are β estimates and 95% credible intervals from univariate and multivariate phylogenetic Bayesian ordinal mixed-effect models assessing potential drivers of sexual selection in birds, restricted to species with highest data quality (scored 3 or 4; *n* = 7592 species). Models were run on a sample of 50 phylogenetic trees extracted from www.birdtree.org (Jetz *et al*. 2012), grafted to the Prum *et al*. (2015) genomic backbone. See Supplementary methods for detailed model descriptions.

| Predictor | Univariate |  | Multivariate |  |
| --- | --- | --- | --- | --- |
|  | Estimate | 95% CI | Estimate | 95% CI |
| 1<sup>ry</sup> consumer | 0.48 | -0.01, 0.98 | 0.62 | 0.10, 1.16 |
| Migration | 1.87 | 1.46, 2.31 | 0.50 | 0.03, 0.96 |
| Territoriality | -3.03 | -3.66, -2.43 | -3.17 | -3.83, -2.55 |
| Seasonality | 2.45 | 2.04, 2.90 | 2.11 | 1.68, 2.58 |
| 1<sup>ry</sup> consumer × territoriality | - | - | 0.74 | -0.12, 1.59 |
| 1<sup>ry</sup> consumer × seasonality | - | - | -1.14 | -1.85, -0.44 |

**Table S6.** Collinearity between multivariate model predictors. Results shown are Variance Inflation Factors (VIF) calculated for multivariate models predicting sexual selection in birds. The two columns show results for all birds (*n* = 9836 species) and a subset of species with high certainty data (*n* = 7592 species). High certainty data was defined as species scored 3 or 4 for data certainty (see Extended Data Table 2). All VIF were below 3, suggesting that collinearity should not impact model interpretation.

| Predictor | Certainty 1-4 | Certainty 3-4 |
| --- | --- | --- |
| 1<sup>ry</sup> consumer | 1.21 | 1.23 |
| Migration | 1.32 | 1.35 |
| Territoriality | 1.24 | 1.24 |
| Seasonality | 1.27 | 1.30 |
| 1<sup>ry</sup> consumer × territoriality | 1.20 | 1.17 |
| 1<sup>ry</sup> consumer × seasonality | 1.05 | 1.06 |

#### Supplementary figures

**Figure S1.**
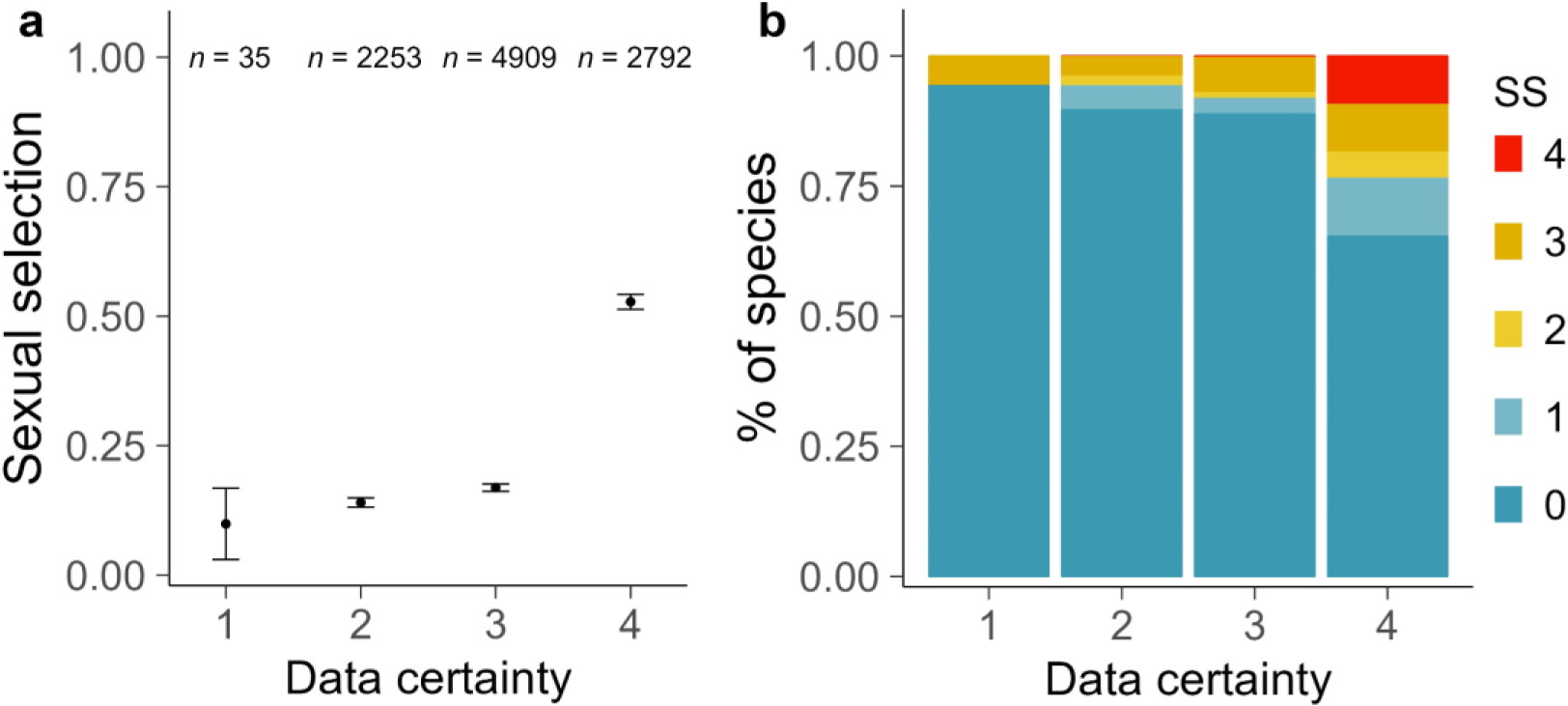
Sexual selection strength across data certainty partitions. Plots show how the certainty in data for each species affects (**a**) average sexual selection and (**b**) variation in sexual selection score. In (**a**) and (**b**), sexual selection is scored from 0 (strict monogamy) to 4 (extreme polygamy; see Extended Data Table 1) and data certainty is scored from 1 (no direct or indirect evidence) to 4 (direct evidence published in primary and secondary literature; see Extended Data Table 2). In (**a**), points show average sexual selection for each category of data certainty; whiskers denote one standard error; sample sizes are the total number of species in each data certainty partition. In (**b**), stacked bars show the relative proportion of sexual selection scores across each category of data certainty. Bars are coloured according to sexual selection score (SS) (blue = 0; red = 4).

**Figure S2.**
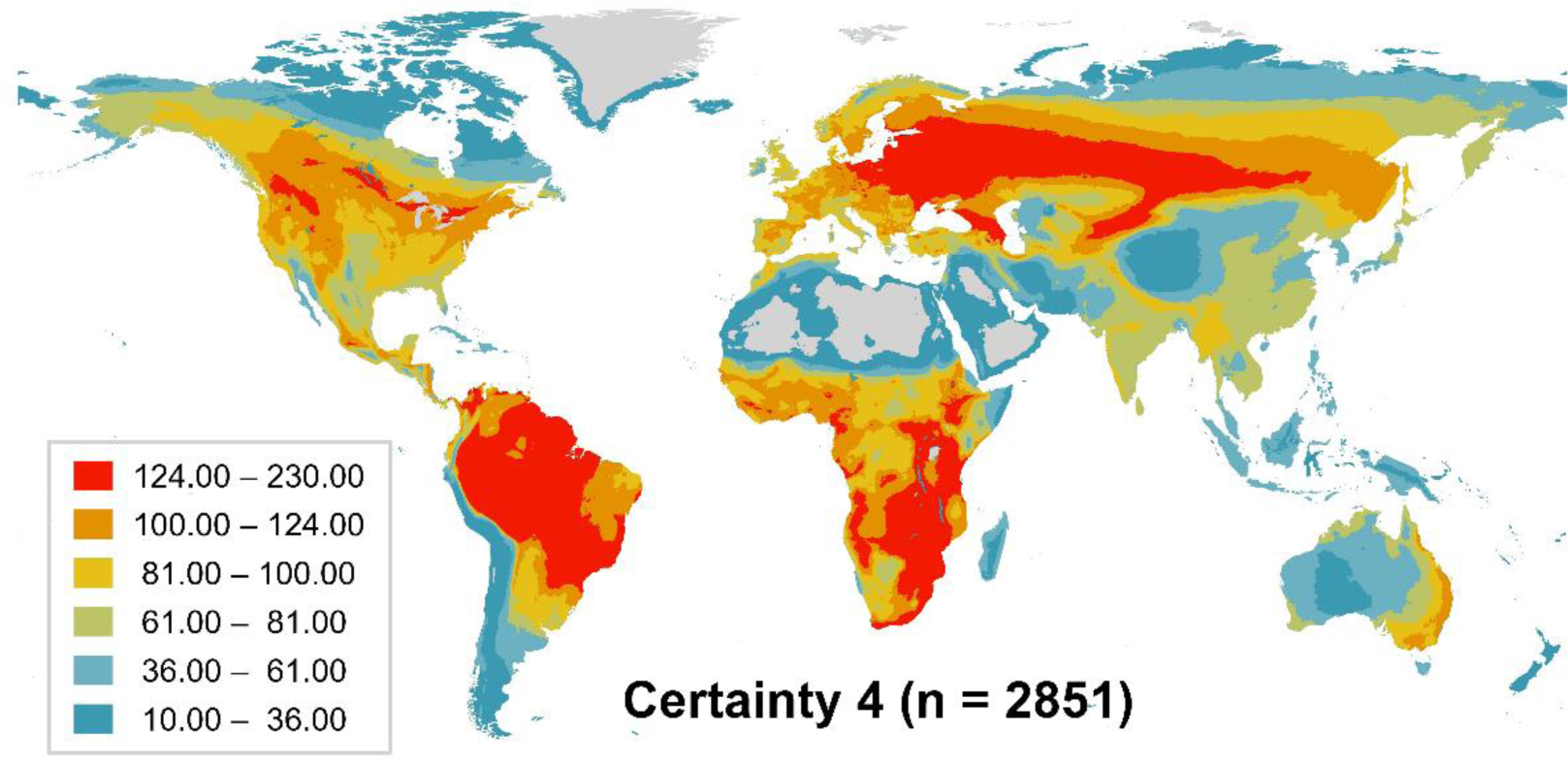
The geographical distribution of species with high-certainty data. Worldwide species richness for 2851 bird species with high data certainty (score = 4) included in a global phylogeny (www.birdtree.org; Jetz *et al*. 2012). Species were scored as 4 based on direct evidence published in primary and secondary literature (see Extended Data Table 2). Cell values represent the total number of high-certainty species, calculated by counting the number of geographical range maps overlapping more than 50% of each 5-km grid cell. Cell values were grouped into equal-sized bins to reduce the skew in the colour scale caused by outlier cells with high species richness. Cells with <10 species were excluded.

